# CLAMP and Zelda function together to promote *Drosophila* zygotic genome activation

**DOI:** 10.1101/2020.07.15.205054

**Authors:** Jingyue Ellie Duan, Leila E. Rieder, Megan M. Colonnetta, Annie Huang, Mary McKenney, Scott Watters, Girish Deshpande, William T. Jordan, Nicolas L. Fawzi, Erica N. Larschan

## Abstract

During the essential and conserved process of zygotic genome activation (ZGA), chromatin accessibility must increase to promote transcription. *Drosophila* is a well-established model for defining mechanisms that drive ZGA. Zelda (ZLD) is a key pioneer transcription factor (TF) that promotes ZGA in the *Drosophila* embryo. However, many genomic loci that contain GA-rich motifs become accessible during ZGA independent of ZLD. Therefore, we hypothesized that other early TFs that function with ZLD have not yet been identified, especially those that are capable of binding to GA-rich motifs such as CLAMP. Here, we demonstrate that *Drosophila* embryonic development requires maternal CLAMP to: 1) activate zygotic transcription; 2) increase chromatin accessibility at promoters of specific genes that often encode other essential TFs; 3) enhance chromatin accessibility to facilitate ZLD occupancy at a subset of key embryonic promoters. Thus, maternal CLAMP functions with ZLD in a pioneer-like role to drive zygotic genome activation.

## INTRODUCTION

During zygotic genome activation (ZGA), dramatic reprogramming occurs in the zygotic nucleus to initiate global transcription and prepare the embryo for further development (Jukam et al., 2017). Chromatin changes that activate the zygotic genome during ZGA rely on cooperation among transcription factors (TFs) (Lee et al., 2014). However, only pioneer TFs (Cirillo and Zaret, 1999; Mayran and Drouin, 2018) can bind to closed chromatin before ZGA because most TFs cannot bind to nucleosomal DNA (Soufi et al., 2015).

In *Drosophila,* the pioneer TF Zelda (ZLD; Zinc-finger early *Drosophila* activator) plays a key role during ZGA (Liang et al., 2008). ZLD exhibits several critical characteristics of pioneer TFs, including: 1) binding to nucleosomal DNA (Sun et al., 2015; McDaniel et al., 2019); 2) regulating transcription of early zygotic genes (Harrison et al., 2011); and 3) modulating chromatin accessibility to increase the ability of other non-pioneer TFs to bind to DNA (Schulz et al., 2015). However, a large subset of ZLD binding sites (60%) are highly enriched for GA-rich motifs and have constitutively open chromatin even in the absence of ZLD (Schulz et al., 2015). Therefore, we and others (Schulz et al., 2015) hypothesized that other pioneer TFs which directly bind to GA-rich motifs work together with ZLD to activate the zygotic genome.

GAGA-associated Factor (GAF, Farkas et al., 1994) and Chromatin-Linked Adaptor for Male-Specific Lethal (MSL) Proteins (CLAMP, Soruco et al., 2013) are two of few known TFs that can bind to GA-rich motifs and regulate transcriptional activation in *Drosophila* (Fuda et al., 2015; Kaye et al., 2018). GAF performs several essential functions in early embryos, including chromatin remodeling (Shimojima et al., 2003; Judd et al., 2020; Gaskill et al., 2021) and RNA Pol II recruitment (Li et al., 2013; Fuda et al., 2015; Duarte et al., 2016), and is required for embryonic nuclear divisions (Bhat et al., 1996).

CLAMP is a GA-binding TF essential for early embryonic development (Rieder et al., 2017) that binds to promoters and plays several vital roles including opening chromatin on the male X-chromosome to recruit the MSL dosage compensation complex (J. Urban et al., 2017; Rieder et al., 2019) and activating coordinated regulation of the histone genes at the histone locus (Rieder et al., 2017). Therefore, we hypothesized that CLAMP functions with ZLD as a pioneer-like factor to promote zygotic genome activation.

Here, we first demonstrate that depleting maternal CLAMP disrupts transcription of critical early zygotic genes causing significant phenotypic changes in early embryos. Next, we define several mechanisms by which CLAMP regulates zygotic genome activation: 1) CLAMP activates zygotic transcription via direct binding to target genes; 2) CLAMP binds directly to nucleosomal DNA and increases chromatin accessibility of promoters of a subset of genes that often encode other essential TFs; 3) CLAMP and ZLD regulate each other’s occupancy at promoters. Overall, we determine that CLAMP is an essential pioneer-like factor that functions with ZLD to regulate zygotic genome activation.

## RESULTS

### Depletion of maternal CLAMP disrupts expression of genes that regulate zygotic patterning and cytoskeletal organization in blastoderm embryos

We previously reported that nearly 100% (99.87%) of maternal *clamp* RNAi embryos never hatch and die at early embryonic stages (Rieder et al., 2017), demonstrating that maternally-deposited CLAMP is critical for embryonic development. To assess embryonic phenotypic patterning after maternal *clamp* depletion, we first identified three key early zygotic genes (*even-skipped*, *runt,* and *neurotactin,* **Figure 1-figure supplement 1A**) that have significantly (adjusted *p*-value < 0.05, DESeq2, RNA-seq) reduced expression in early embryos when maternal CLAMP is depleted (Rieder et al., 2017). We then used single-molecule fluorescent in situ hybridization (smFISH) for *even-skipped* and *runt,* and immunostaining for Neurotactin to determine how the depletion of maternal *clamp* or *zld* alters phenotypic patterning and cytoskeletal integrity in blastoderm stage embryos **(Figure 1)**. We validated knockdown of *clamp* and *zld* in early embryos by qRT-PCR and western blotting (**Figure 1-figure supplement 1B, C** and **Figure 1 Source Data 1-2**).

**Figure 1.**
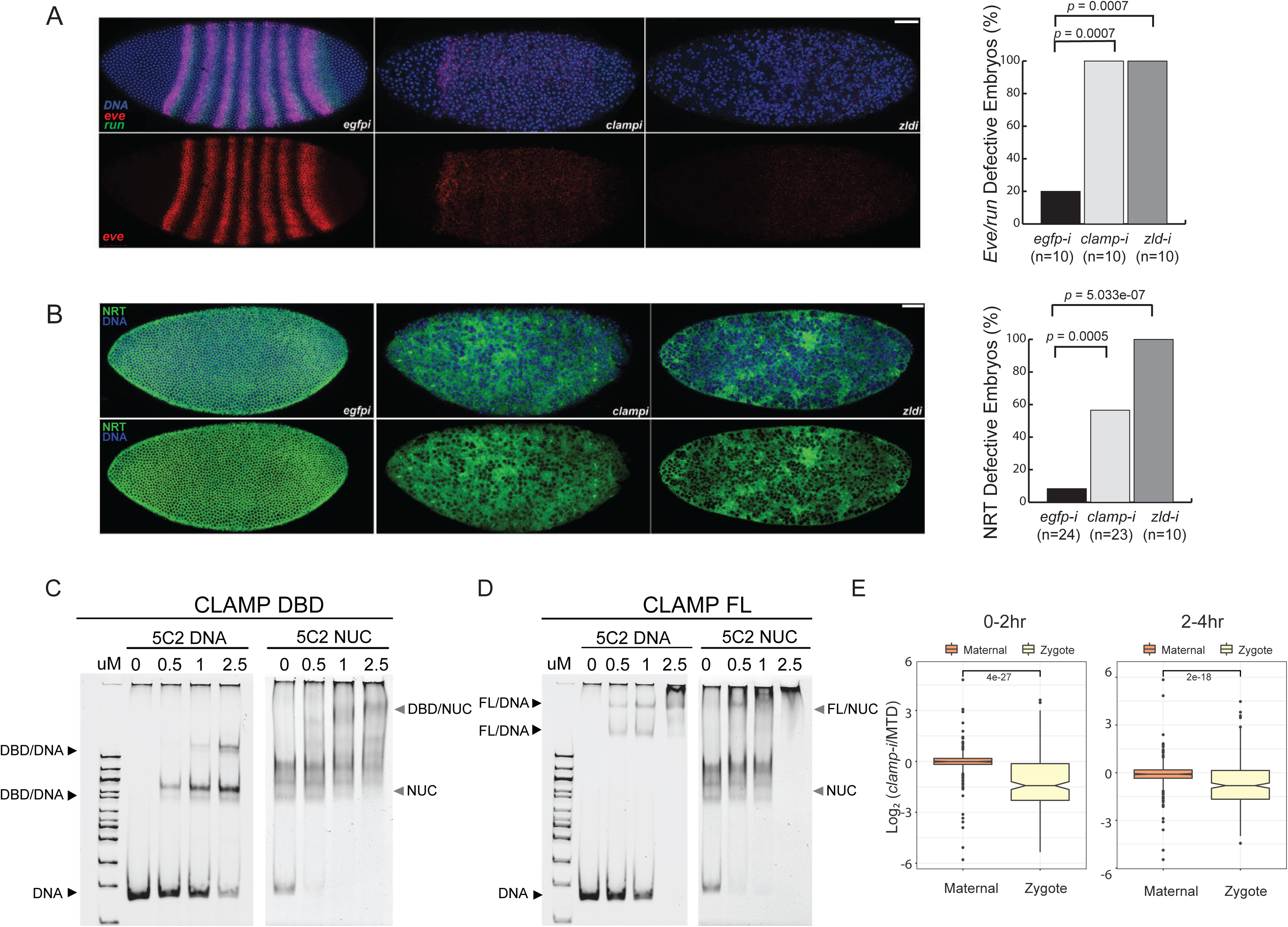
Novel pioneer factor CLAMP is essential for early embryonic development. A. Control maternal *egfp* depletion (left), maternal *clamp* depletion (middle) and maternal *zld* depletion (right) syncytial blastoderm stage embryos probed using smFISH for the pair-rule patterning genes *run* (green) and *eve* (red). Embryos were co-labeled with Hoechst (blue) to visualize nuclei. Scale bar represents 10 µm. Quantification (%) of *eve/run* defective embryos is on the right, *p-value*s were calculated with the *Fisher’s* Exact test; number of embryos is in parentheses. B. Control maternal *egfp* depletion (left), maternal *clamp* depletion (middle) and maternal *zld* depletion (right) syncytial blastoderm stage embryos were assessed for integrity of the developing cytoskeleton using anti-NRT antibody (green) and Hoechst (blue) to label nuclei. Scale bar represents 10 µm. Quantification (%) of NRT defective embryos is on the right, *p-values* were calculated with the *Fisher’s* Exact test; number of embryos is in parentheses. C. Electrophoretic mobility shift assay (EMSA) showing the binding of increasing amounts of CLAMP DNA-binding domain (DBD) fused to MBP to 5C2 naked DNA or 5C2 *in vitro* reconstituted nucleosomes (NUC). Concentrations (uM) of CLAMP DBD increase from left to right. D. EMSA showing the binding of increasing amounts of full-length (FL) CLAMP (fused to MBP) to 5C2 DNA or 5C2 nucleosomes (NUC). Concentrations (uM) of CLAMP FL increase from left to right. E. Effect of maternal *clamp* RNAi on maternally-deposited (orange) or zygotically-transcribed (yellow) gene expression log2 (*clamp*-i/MTD) in 0-2hr (left) or 2-4hr (right). Maternal vs. zygotic gene categories were as defined in Lott et al. (2011). *p*-values of significant expression changes between maternal and zygotic genes were calculated by Mann-Whitney U-test and noted on the plot.

Both *even-skipped (eve)* and *runt (run)* play an important role in embryonic segmentation (Manoukian and Krause, 1992). *Eve* also establishes sharp boundaries between parasegments (Fujioka et al., 1995). Strikingly, when maternally-deposited *clamp* is depleted, we observed the complete disruption of classic seven stripe pair-rule gene expression patterns using smFISH **(Figure 1A, middle)**. Additionally, the nuclei in the embryonic syncytium were disassociated compared to control *egfp* RNAi embryos **(Figure 1A, left)**. Furthermore, the expression of *eve* and *run* was significantly reduced in *clamp* maternal depletion embryos that also failed to form sharp stripe boundaries. We observed similar, but slightly stronger, phenotypic changes in *zld* maternal depletion embryos **(Figure 1A, right)**, indicating that CLAMP and ZLD have critical roles in establishing embryonic patterning in pre-cellular blastoderm embryos. Moreover, all of embryos depleted for maternal *clamp* or maternal *zld* (n = 10) show defective *eve/run* localization (*p* < 0.05, *Fisher’s* exact test).

Next, we used immunostaining to examine the localization of Neurotactin (NRT), a cell adhesion glycoprotein that is expressed early during *Drosophila* embryonic cellularization in a lattice surrounding syncytial blastoderm nuclei (Hortsch et al., 1990). In *clamp* maternal depletion embryos **(Figure 1B, middle)**, we observed dramatically disrupted cellularization and reduced NRT levels. These embryos fail to form the wild type pattern of cytoskeletal elements, which can be seen in the e*gfp* RNAi control embryos **(Figure 1B, left)**. Embryos depleted for maternal *zld* also reveal similar patterns of discordant nuclei. More than 50% of embryos depleted for maternal *clamp* (n = 23) and 100% of embryos depleted for maternal *zld* (n=10) show NRT disruption **(Figure 1B, right)**, (*p* < 0.05, *Fisher’s* exact test). Overall, smFISH and immunostaining results suggest that both maternally-deposited CLAMP and ZLD are essential for early embryonic patterning and development.

### CLAMP binds to nucleosomal DNA *in vivo* and *in vitro*

One of the intrinsic characteristics of pioneer transcription factors is their capacity to bind nucleosomal DNA and compacted chromatin (Cirillo and Zaret, 1999). To test the hypothesis that CLAMP is a pioneer-like TF, we performed electromobility shift assays (EMSAs) that test the intrinsic capability of CLAMP to directly interact with nucleosomes *in vitro*. First, we identified a 240 bp region of the X-linked 5C2 locus **(Figure 1 – figure supplement 1D)** that CLAMP binds to in cultured S2 cells and exhibited decreased chromatin accessibility in the absence of CLAMP (J. Urban et al., 2017). This region is also occupied by a nucleosome **(Figure 1 – figure supplement 1D)**, suggesting that CLAMP promotes accessibility of this region while binding to nucleosomes.

We then performed *in vitro* nucleosome assembly using 240 bp of DNA from the 5C2 locus that contains three CLAMP-binding motifs, and we used 5C2 naked DNA as a control. We found that both the CLAMP DNA binding domain (DBD, **Figure 1C** and **Figure 1 Source Data 3**) and full-length protein (FL, **Figure 1D** and **Figure 1 Source Data 4**) can bind and shift both 5C2 naked DNA and nucleosomes assembled with 5C2 DNA. Increased protein concentration results in a secondary “super” shift species **(Figure 1C, 1D** and **Figure 1 Source Data 3-4)**, indicating that multiple CLAMP molecules may occupy the three CLAMP-binding motifs. Both full length CLAMP and CLAMP DBD are fused to the maltose binding protein (MBP), which we previously demonstrated does not bind to DNA independent of CLAMP or alter the specificity of CLAMP binding (Kaye et al., 2018). Previously, we determined that CLAMP binds specifically to GAGA repeats *in vivo* and *in vitro* (Soruco et al., 2013; Kaye et al., 2018). Here we further demonstrate that the zinc-finger protein CLAMP can directly bind to nucleosomal DNA and generates multiple shift species consistent with the potential to bind to multiple binding sites simultaneously.

### CLAMP regulates zygotic genome activation (ZGA)

To define how CLAMP regulates early embryonic patterning, we examined the effect of maternal CLAMP depletion on the expression of maternally-deposited or zygotically-transcribed genes (Lott et al., 2011) using mRNA-seq data (Rieder et al., 2017). We found that the expression levels of zygotically-transcribed genes but not maternally-deposited genes were significantly downregulated in embryos lacking CLAMP (*p* < 0.001, Mann-Whitney U-test) **(Figure 1E).** Therefore, CLAMP has a specific effect on transcription of zygotic genes similar to that which has been previously reported for ZLD (Liang et al., 2008; Harrison et al., 2011; McDaniel et al., 2019) and confirmed in this study **(Figure 1 – figure supplement 1E)** using stage 5 embryos lacking maternal ZLD (GSE65837, Schulz et al., 2015).

### CLAMP regulates chromatin accessibility in early embryos

An essential characteristic of pioneer transcription factors is that they can establish and maintain the accessibility of their DNA target sites, allowing other TFs to bind to DNA and activate transcription (Zaret and Carroll, 2011; Iwafuchi-Doi et al., 2016). We previously used MNase-seq (J. Urban et al., 2017) to determine that CLAMP guides MSL complex to GA-rich sequences by promoting an accessible chromatin environment on the male X-chromosome in cultured S2 cells. Furthermore, GA-rich motifs are enriched in regions that remain accessible in the absence of the pioneer factor ZLD (Schulz et al., 2015). Therefore, we hypothesized that CLAMP regulates chromatin accessibility at some ZLD-independent GA-rich loci during ZGA.

To test our hypothesis, we performed Assay for Transposase-Accessible Chromatin using sequencing (ATAC-seq) on 0-2hr (pre-ZGA) and 2-4hr (post-ZGA) embryos with wild-type levels of CLAMP [maternal triple GAL4 driver (MTD) alone (Ni et al., 2011)] and embryos depleted for maternal CLAMP using RNAi driven by the MTD driver (*clamp-i*). We identified differentially accessible (DA) regions **(Figure 2A** and **Figure 2-√ table 1)** by comparing ATAC-seq reads between MTD and *clamp-i* embryos using Diffbind (Stark and Brown, 2019). PCA plots (**Figure 2-figure supplement 1A & D)** show that the first principal component (PC) explains 86%-87% of the variation between MTD and *clamp-i* embryos. However, we also observed that 5-6% of the variation among sample replicates is explained by PC2, suggesting the presence of some developmental diversity within sample groups.

**Figure 2.**
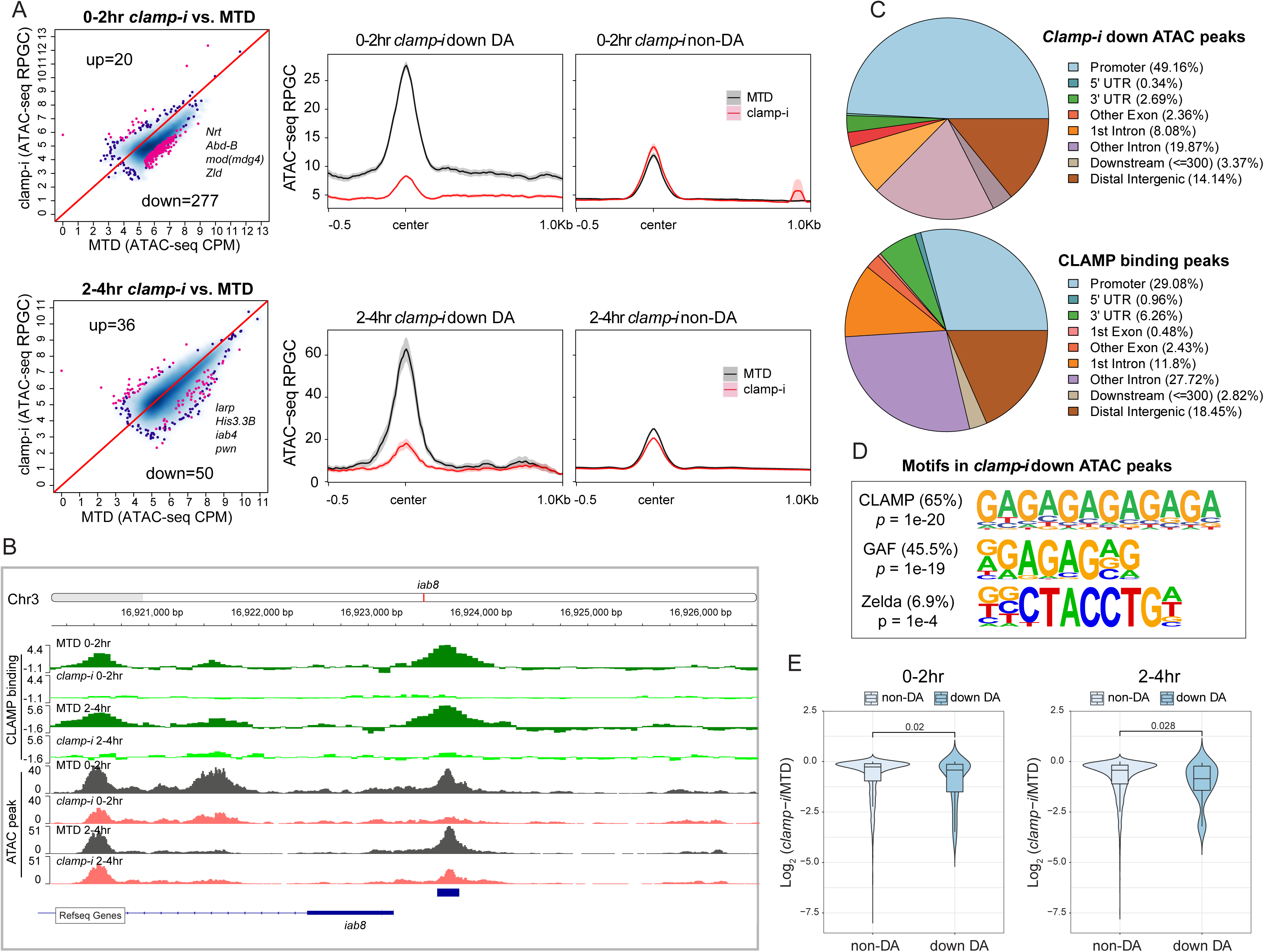
CLAMP regulates chromatin accessibility of a subset early zygotic genome. A. Differential accessibility (DA) analysis (left) of ATAC-seq from MTD embryos versus *clamp-i* embryos in 0-2hr or 2-4hr. Blue dots indicate non-differentially accessible sites (non-DA). Pink dots indicate significant (FDR < 0.1) differential peaks after maternal *clamp* RNAi, identified by DiffBind (DESeq2) (DA peaks). The number of peaks and representative genes in each class is noted on the plot. Average ATAC-seq signal (right) in reads per genome coverage (RPGC) 1x normalization in 0-2hr or 2-4hr embryos after maternal *clamp* RNAi centered on open chromatin (<= 100bp) peaks identified significant changes upon maternal *clamp* RNAi. B. Example of IGV views of genomic locus *iab-8* bound by CLAMP (ChIP-seq) which shows significantly decreased CLAMP binding and ATAC-seq signal after *clamp* RNAi. C. Genomic features of regions that require CLAMP for chromatin accessibility (ATAC-seq) compared with all CLAMP binding sites (ChIP-seq). D. Top motifs enriched in regions that require CLAMP for chromatin accessibility. Enrichment *p*-value and percentage of sequences are noted. E. Violin plot comparing gene expression in CLAMP-mediated changes and unchanged differential accessibility regions in 0-2hr or 2-4hr embryos after maternal *clamp* RNAi. *p*-values of significant expression changes of CLAMP down-DA and non-DA were calculated by Mann-Whitney U-test and noted on the plot.

Despite some variation among replicates, the high Pearson correlation for DA regions between replicates indicates robust reproducibility of these sites **(Figure 2-figure supplement 1B & E).** We identified a subset of genomic regions that exhibit significantly reduced chromatin accessibility in the absence of CLAMP (**Figure 2A**, 0-2hr: 277; 2-4hr: 50 and **Figure 2-supplementary table 1**), indicating that chromatin accessibility of these genomic loci (DA sites) requires CLAMP. Moreover, DA sites include promoters of many genes essential for early embryogenesis such as *Nrt, Abd-B, mod(mdg4),* and *zld,* which encodes the ZLD TF (**Figure 2A)**. A smaller number of loci (0-2hr: 20; 2-4hr: 36) increased their accessibility in the absence of CLAMP **(Figure 2A)**. Gene Ontology (GO) analysis **(Figure 2-figure supplement 1C & F)** indicates that CLAMP increases accessibility of chromatin regions that are mainly within DNA-binding, RNA Pol II binding, and enhancer-binding TF encoding genes **(Figure 2-figure supplement 1C & F)**. While CLAMP strongly regulates chromatin accessibility non-redundantly with other factors at a subset of genomic loci, CLAMP target genes are key for early development, consistent with the dramatic patterning defects observed after depleting maternal CLAMP **(Figure 1A)**.

Furthermore, a subset [26.7% (74/277) at 0-2hrs; 90% (45/50) at 2-4hrs] of DA regions are directly bound by CLAMP, suggesting that CLAMP directly regulates their chromatin accessibility. For example, the *iab8* promoter, which is located within the essential *Drosophila Hox* cluster that controls body plan patterning, is directly bound by CLAMP and shows a reduction in chromatin accessibility after *clamp* RNAi **(Figure 2B)**. We also defined the distribution of DA sites and CLAMP binding sites throughout the genome **(Figure 2C)**. While DA sites were significantly (*p* < 0.05, *Fisher’s* exact test) enriched at promoter regions (49.16%), CLAMP binds almost equally frequently to both promoters (29.08%) and introns that are not first introns (27.72%). Therefore, CLAMP is required to establish or maintain open chromatin largely at promoters, but may also play other roles at intronic regions. Furthermore, motif analysis identified both GA-rich motifs and ZLD motifs enriched at regions that require CLAMP for their accessibility in 0-2hr embryos **(Figure 2D)**. These data suggest that CLAMP may also regulate the accessibility of some ZLD binding sites, a hypothesis that we discuss further below.

We next determined whether CLAMP-mediated chromatin accessibility could specifically drive early transcription by examining the relationship between the chromatin accessibility (DA, ATAC-seq) changes and gene expression of the nearest gene as measured by RNA-seq (Rieder et al., 2017). We observed significant (*p*-values < 0.05, Mann-Whitney U-test) reduction in expression after maternal CLAMP depletion of genes at which CLAMP mediates chromatin accessibility (DA genes) compared with genes at which CLAMP does not regulate chromatin accessibility (non-DA genes) (**Figure 2E)**. Overall, our results indicate that CLAMP promotes chromatin accessibility and transcription of a subset of other essential TF genes during ZGA, which is consistent with the extensive developmental defects caused by maternal CLAMP depletion.

### CLAMP and ZLD regulate each other’s binding to a subset of promoters

To directly determine how CLAMP and ZLD impact each other’s binding, we performed ChIP-seq for CLAMP and ZLD in control MTD embryos and embryos that were maternally depleted for each factor with RNAi at the same two time points we used for our ATAC-seq experiments: before ZGA (0-2hr) and during and after ZGA (2-4hr) **(Figure 3A-B, Figure 3-figure supplement 1A-B** and **Table 1)**. Overall, there are more ZLD peaks (0-2hr: 6,974; 2-4hr: 8,035) across the whole genome than CLAMP peaks (0-2hr: 4,962, 2-4hr: 7,564) in control MTD embryos. As we hypothesized, CLAMP and ZLD peaks significantly overlap (*p* < 0.05, Hypergeometric test, N=15,682 total fly genes) **(Figure 3-figure supplement 1C)**.

**Figure 3.**
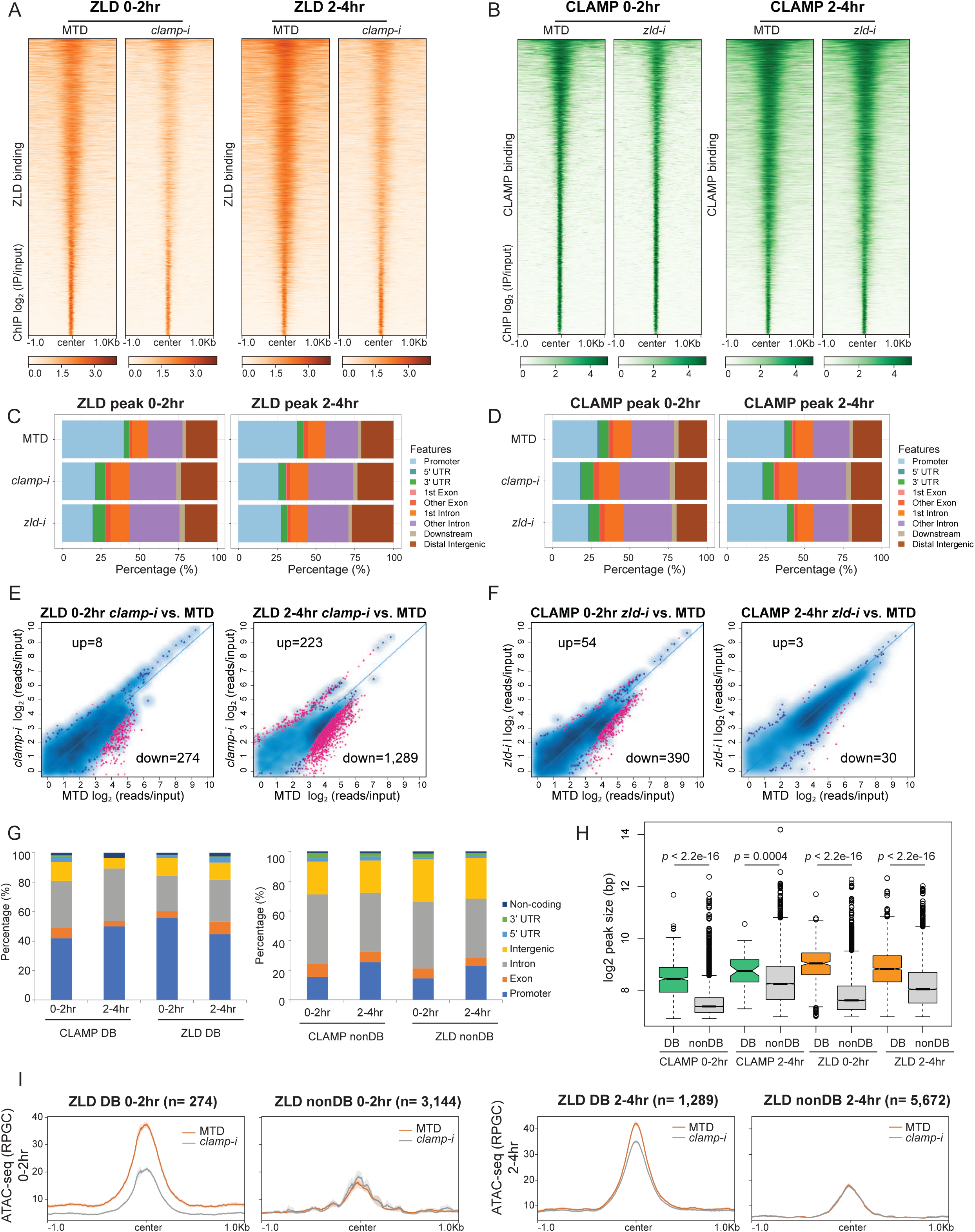
CLAMP and ZLD depend on each other for binding at a subset of sites. A. ZLD peaks in 0-2hr and 2-4hr MTD and maternal *clamp* RNAi embryos. Data is displayed as a heatmap of *z*-score normalized ChIP-seq (log2 IP/input) reads in a 2 Kb region centered at each peak. Peaks in each class are arranged in order of decreasing *z*-scores in control MTD embryos. B. CLAMP peaks in 0-2hr and 2-4hr MTD and maternal *zld* RNAi embryos. Data is displayed as a heatmap of z-score normalized ChIP-seq (log2 IP/input) reads in a 2 Kb region centered around each peak. Peaks in each class are arranged in order of decreasing z-scores in control MTD embryos. C. Genomic distribution fractions in promoters, UTRs, exon, intron and intergenic regions for ZLD peaks in the *Drosophila* genome (dm6) in 0-2hr and 2-4hr embryos (MTD, *clamp*-i and *zld*-i). D. Genomic distribution fractions in promoters, UTRs, exon, intron and intergenic regions for CLAMP peaks in the *Drosophila* genome (dm6) in 0-2hr and 2-4hr embryos (MTD, *clamp*-i and *zld*-i). E. Differential binding (DB) analysis of ZLD ChIP-seq. Mean difference (MA) plots of ZLD peaks in MTD embryos versus *clamp*-i embryos in 0-2hr (left) or 2-4hr (right). Blue dots indicate non-differential binding sites. Pink dots indicate significant (FDR < 0.05) differential peaks identified by DiffBind (DESeq2). The number of peaks changed in each direction is noted on the plot. F. Differential binding (DB) analysis of CLAMP ChIP-seq. MA plots of CLAMP peaks from MTD embryos versus *zld*-i embryos in 0-2hr (left) or 2-4hr (right). Blue dots indicate non-differential binding (non-DB) sites. Pink dots indicate significant (FDR < 0.05) differential binding (DB) peaks identified by DiffBind (DESeq2). Number of peaks in each direction is noted on the plot. G. Stacked bar plots of CLAMP and ZLD down-DB (left) and CLAMP and ZLD non-DB peaks (right) distribution fraction in the *Drosophila* genome (dm6) in 0-2hr and 2-4hr embryos. H. Box plot of the peak sizes in CLAMP and ZLD down-DB and non-DB peaks in 0-2hr and 2-4hr embryos. *p*-values of significant size difference between down-DB and non-DB peaks were calculated by Mann-Whitney U-test and noted on the plot. I. Average profiles of ATAC-seq signal coverage show chromatin accessibility at ZLD down-DB (orange line) and non-DB (grey line) sites in 0-2hr (left panel) or 2-4hr (right panel) MTD and *clamp*-i embryos. Number of peaks is noted on the plot.

**Table 1.** The number of total and differentially bound peaks for CLAMP and ZLD in control MTD, *clamp-i* and *zld-i* embryos.

| ChIP-seq Peaks | CLAMP |  |  | ZLD |  |  |
| --- | --- | --- | --- | --- | --- | --- |
|  | *MTD* | *clamp-i* | *zld-i* | *MTD* | *clamp-i* | *zld-i* |
| 0-2hr | 4,962 | 3,488 | 4,746 | 6,974 | 3,687 | 4,650 |
| 2-4hr | 7,564 | 4,064 | 8,279 | 8,035 | 4,687 | 6,420 |
| Differential Binding (DiffBind, DEseq2) | *MTD vs. zld-i* |  |  | *MTD vs. clamp-i* |  |  |
|  | up-DB | down-DB | non-DB | up-DB | down-DB | non-DB |
| 0-2hr | 54 | 390 | 4,184 | 8 | 274 | 3,144 |
| 2-4hr | 3 | 30 | 7,351 | 223 | 1,289 | 5,672 |

Before ZGA (0-2hrs), ZLD showed higher enrichment at promoters than CLAMP. In contrast, CLAMP had a similar distribution to ZLD at the 2-4hr time point, indicating that CLAMP binds to promoters later than ZLD **(Figure 3C-D)**. Depletion of either maternal *zld* or *clamp* mRNA altered the genomic distribution of CLAMP and ZLD: both factors shifted their occupancy from promoters to introns. Interestingly, maternal *zld* RNAi no longer affects CLAMP binding to promoters at the 2-4hr time point, which may be because the maternal RNAi is not as effective at reducing ZLD protein levels at this time point **(Figure 3-figure supplement 1B)**. Therefore, CLAMP and ZLD change the distribution of each other’s occupancy to increase their binding to promoters instead of introns.

Next, we defined the sites that showed differential binding (DB, **Figures 3E-F** and **Table 1**) of CLAMP and ZLD in the absence of each other’s maternally-deposited mRNA using Diffbind (Stark and Brown, 2019). We found a significant reduction of ZLD binding in the absence of CLAMP: there were 274 (0-2hr) and 1,289 (2-4hr) sites where ZLD binding decreased in *clamp-i* embryos compared to MTD controls (down-DB) **(Figures 3E, Figure 3-figure supplement 1D**, and **Table 1)**. Fewer ZLD binding sites increased in occupancy after *clamp* RNAi: 8 (0-2hr) and 233 (2-4hr) sites (up-DB). In contrast, loss of ZLD had a weaker impact on CLAMP binding: 390 (0-2 hr) and 30 (2-4 hr) down-DB sites **(Figures 3F, Figure 3-figure supplement 1E**, and **Table 1)**. We identified very few sites where CLAMP occupancy increases after *zld* RNAi (up-DB sites: 0-2hr: 54, 2-4hr: 3).

Moreover, the CLAMP down-DB sites and the ZLD down-DB sites significantly overlap with each other (*p* < 0.05, Hypergeometric test, N=15,682 total fly genes) at both timepoints **(Figure 3-figure supplement 1F)**. For example, *iab-8*, an essential *Hox* cluster gene at which CLAMP regulates chromatin accessibility (DA), is also one of the 95 genomic loci at which CLAMP and ZLD promote each other’s occupancy **(Figure 3-figure supplement 1F-G)**.

To further understand how CLAMP and ZLD bind to dependent (down-DB) and independent (non-DB) sites, we determined the genomic distribution of these two types of sites. Overall, dependent peaks are much broader in size and are located at promoters **(Figure 3G, Figure 3-figure supplement 1H** and **Figure 3-figure supplement 2A)**. In contrast, independent sites are narrower and located within introns **(Figure 3G, Figure 3-figure supplement 1H** and **Figure 3-figure supplement 2A)**. On average, the peak size of dependent sites (down-DB: 400-500bp) is almost double that of independent sites (non-DB: 200-250bp) with significant differences in peak size for both TFs at both time points (*p* < 0.001, Mann-Whitney U-test) **(Figures 3H)**.

Previous proteomic studies (J. A. Urban et al., 2017; Hamm et al., 2017, 2015) found no evidence that CLAMP and ZLD directly contact each other at the protein level, suggesting that CLAMP and ZLD regulate each other via binding to their DNA motifs. Therefore, we analyzed the motifs enriched at dependent (down-DB) and independent (non-DB) sites. We found that dependent sites are enriched for motifs specific for the required protein, which are not present at the independent sites **(Figure 3-figure supplement 2B).** For example, the ZLD motif is only enriched at sites where CLAMP requires ZLD for binding (CLAMP down-DB) but not at sites where CLAMP binds independently of ZLD (CLAMP non-DB). Similarly, the CLAMP motif is only enriched at sites where ZLD requires CLAMP for binding (ZLD down-DB) **(Figure 3-figure supplement 2B).** Therefore, the presence of specific CLAMP and ZLD motifs correlates with their ability to promote each other’s binding.

Given the cooperative relationship between CLAMP and ZLD binding to chromatin, we measured chromatin accessibility changes at their dependent and independent sites that we defined from ChIP-seq data **(Figures 3I** and **Figure 3-figure supplement 2C).** We found the average ATAC-seq signals were significantly reduced at sites where ZLD is dependent on CLAMP to bind (ZLD down-DB sites) in *clamp-i* embryos compared to MTD controls **(Figure 3I)**. Furthermore, the accessibility at sites where ZLD binds independently of CLAMP (ZLD non-DB) is lower than that at ZLD DB sites but remains unchanged upon *clamp* RNA-i **(Figure 3I)**. Therefore, the chromatin accessibility changes we observe over broader regions are enriched at specific loci where CLAMP promotes ZLD binding.

Sites where ZLD regulates CLAMP binding (CLAMP down-DB) have high chromatin accessibility while sites where CLAMP binds independently (CLAMP non-DB) of ZLD showed low chromatin accessibility **(Figure 3-figure supplement 2C)**. Interestingly, accessibility slightly increases upon the loss of ZLD at sites where CLAMP requires ZLD for binding at 0-2hr **(Figure 3-figure supplement 2C).** However, an active TF binding to DNA can prevent Tn5 cleavage at genomic regions (Yan et al., 2020). Therefore, loss of ZLD and CLAMP binding could result in a perceived accessibility increase, as measured by ATAC-seq, which does not necessarily reflect a repressive function for ZLD.

In summary, CLAMP and ZLD increase each other’s occupancy by binding to their motifs and altering chromatin accessibility. These data support a model in which CLAMP and ZLD increase each other’s occupancy at promoters of a subset of genes that often encode other transcription factors.

### CLAMP and ZLD function together to regulate genes during zygotic genome activation

CLAMP and ZLD both specifically regulate zygotic transcription **(Figure 1E)** (Liang et al., 2008; Harrison et al., 2011; McDaniel et al., 2019, Rieder et al., 2017). To further understand how CLAMP and ZLD function to regulate ZGA, we compared the transcriptional roles of CLAMP and ZLD in early embryos at genes that have different temporal expression patterns as defined in Li et al. (2014). First, we found that both CLAMP and ZLD are present at genes expressed throughout ZGA although CLAMP binding is more often present at mid- and late-transcribed zygotic genes (categories defined in Li et al., 2014), while ZLD binding is more often present at early-transcribed zygotic genes **(Figure 4A-B)**.

**Figure 4.**
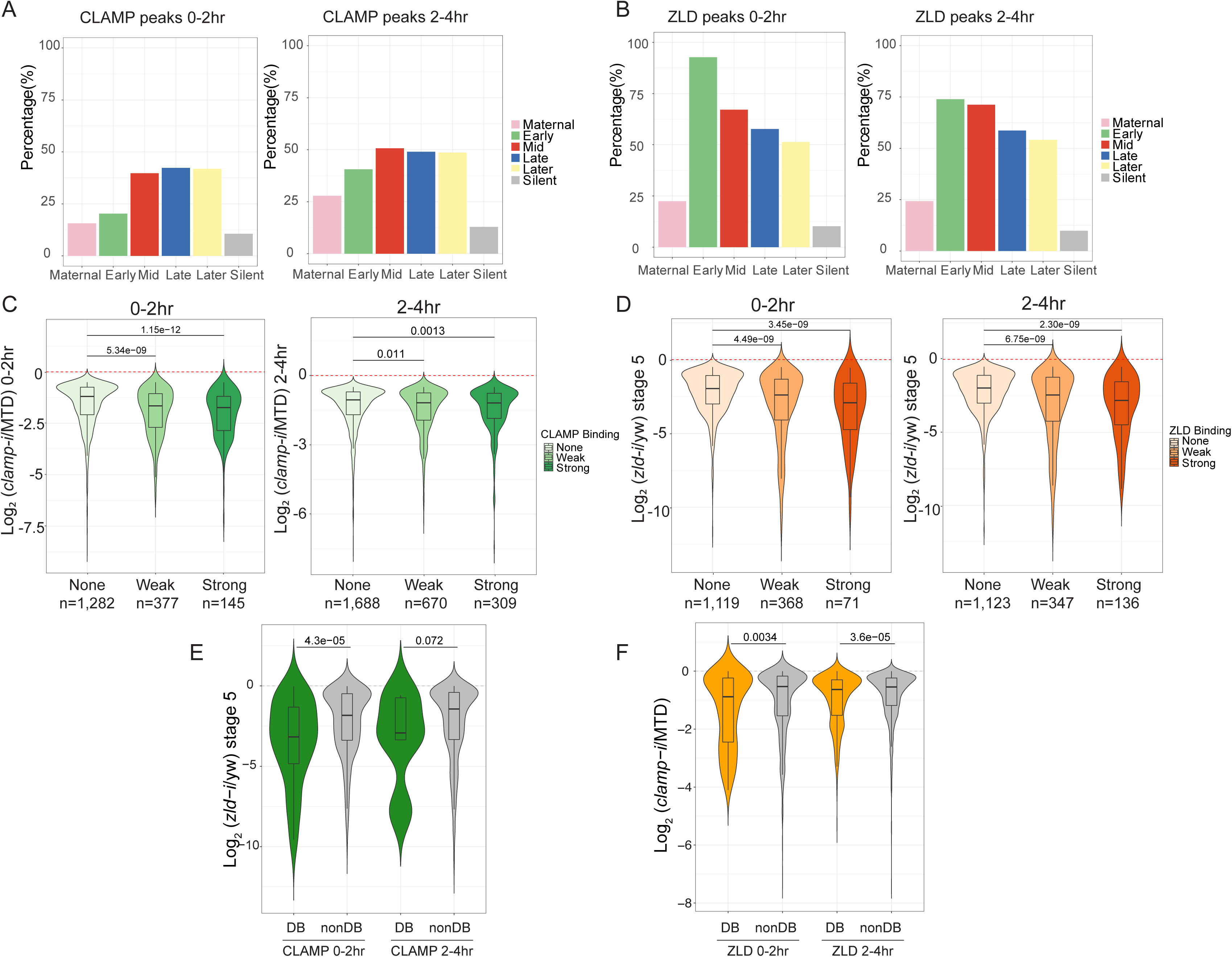
CLAMP and ZLD function together in zygotic genome activation. A. Percentage of CLAMP binding sites in 0-2hr and 2-4hr embryos distributed in maternal (n = 646), early (n = 69), mid-(n = 73), late- (n = 104), later (n = 74), and silent (n = 921) genes (peaks within a 1 Kb promoter region and gene body). Gene categories were defined in Li et al. (2014). B. Percentage of ZLD binding sites in 0-2hr and 2-4hr embryos distributed in maternal (n = 646), early (n = 69), mid-(n = 73), late- (n = 104), later (n = 74), and silent (n = 921) genes (peaks within a 1 Kb promoter region and gene body). Gene categories were defined in Li et al. (2014). C. Gene expression changes caused by maternal *clamp* RNAi (Rieder et al., 2017) at genes with strong, weak, and no CLAMP binding as measured by ChIP-seq in 0-2hr (left) or 2-4hr (right) embryos. *p*-values of significant expression changes of CLAMP bindings were calculated by Mann-Whitney U-test and noted on the plot. D. Gene expression changes caused by maternal *zld* RNAi (Schulz et al., 2015) at genes with strong, weak, and no ZLD binding as measured by ChIP-seq in 0-2hr (left) or 2-4hr (right) embryos. *p*-values of significant expression changes of ZLD bindings were calculated by Mann-Whitney U-test and noted on the plot. E. Gene expression changes caused by maternal *zld* RNAi (Schulz et al., 2015) at genes with CLAMP down-DB and non-DB that defined in wt vs. *zld*-0-2hr and 2-4hr embryos. *p*-values of significant expression changes of CLAMP down-DB and non-DB were calculated by Mann-Whitney U-test and noted on the plot. F. Gene expression changes caused by maternal *clamp* RNAi (Rieder et al., 2017) at genes with ZLD down-DB and non-DB that defined in MTD vs. *clamp*-i 0-2hr and 2-4hr embryos. *p*-values of significant expression changes of ZLD down-DB and non-DB were calculated by Mann-Whitney U-test and noted on the plot.

We next asked whether the ability of CLAMP to bind to genes directly regulates zygotic gene activation by integrating ChIP-seq with RNA-seq data (Schulz et al., 2015; Rieder et al., 2017). We determined that genes strongly bound or weakly bound by CLAMP showed a significant (*p* < 0.001, Mann-Whitney U-test) level of gene expression reduction after *clamp* RNAi (Rieder et al., 2017) compared to unbound genes **(Figure 4C).** We also observed a significant (*p* < 0.001, Mann-Whitney U-test) change in gene expression in maternal *zld-i* embryos (Schulz et al., 2015) for the genes that are strongly bound by ZLD **(Figure 4D)**. Also, the magnitude of the transcriptional changes is similar for genes that are bound by CLAMP or ZLD. Together, these data indicate that CLAMP regulates transcription of zygotic genes by directly binding to target genes.

Next, we tested the hypothesis that CLAMP and ZLD could regulate each other’s binding to precisely drive the transcription of target genes. To test this hypothesis, we plotted the gene expression changes caused by depleting maternal ZLD at genes where CLAMP regulates ZLD binding **(Figure 4E)**. The depletion of maternal *zld* significantly (*p* = 4.3e-5, Mann-Whitney U-test) reduces the expression of genes where ZLD regulates CLAMP binding (down-DB) more than sites where CLAMP binds independently of ZLD (non-DB) at the 0-2hr and 2-4hr (*p* = 0.072) time points **(Figure 4E)**. Therefore, ZLD may specifically regulate zygotic genes at which ZLD promotes CLAMP binding. Also, compared to genes where ZLD binds independent of CLAMP, genes where ZLD binding is regulated by CLAMP had a significant (*p* < 0.001, Mann-Whitney U-test) expression reduction after *clamp* RNAi at both 0-2hr and 2-4hr time points **(Figure 4F)**. Thus, CLAMP may regulate the transcription of genes targeted by ZLD by promoting ZLD binding.

Furthermore, sites where CLAMP and ZLD require each other for binding are enriched for motifs specific for the required protein **(Figure 3-figure supplement 2B)**. Therefore, the presence of specific CLAMP and ZLD motifs correlates with their ability to promote each other’s binding which further regulates the expression of each other’s target genes.

### CLAMP and ZLD regulate gene expression via modulating chromatin accessibility

To determine how direct binding of CLAMP and ZLD relates to zygotic chromatin accessibility, we integrated ChIP-seq and ATAC-seq data. First, we defined four classes of CLAMP-related peaks (DA with CLAMP, DA without CLAMP, Non-DA with CLAMP, Non-DA without CLAMP in **Table 2** and **Table 2 - Source Data 1**). We also obtained ZLD-related ATAC-seq data (Hannon et al., 2017; Soluri et al., 2020) that was generated from embryos laid by wildtype (wt) mothers or mothers with *zld* germline clones (*zld-*) at the nuclear cycle 14 (NC14) +12 min time point and integrated it with ChIP-seq data from the closest time point from this study (0-2hr embryos). In this way, we defined four classes of genomic loci related to ZLD: DA with ZLD, DA without ZLD, Non-DA with ZLD, Non-DA without ZLD **(Table 2** and **Table 2 - Source Data 1)**.

**Table 2.** The number of peaks in 4 types of CLAMP or ZLD mediated regions.

| ATAC-seq peaks (0-2hr) | DA w/<br>CLAMP | DA w/o<br>CLAMP | Non-DA w/<br>CLAMP | Non-DA w/o<br>CLAMP |
| --- | --- | --- | --- | --- |
| 16,597 | 74 | 203 | 1,239 | 15,081 |
| ATAC-seq peaks (Hannon et al., 2017) (NC14+12min) | DA w/ ZLD | DA w/o ZLD | Non-DA w/<br>ZLD | Non-DA w/o<br>ZLD |
| 19,146 | 976 | 2,782 | 2,010 | 13,378 |
| CLAMP ZLD co-bound open chromatin regions | Type I<br>(Both DA) | Type II<br>(CLAMP DA) | Type III<br>(ZLD DA) | Type IV<br>(Both non-DA) |
| 525 | 5 | 23 | 123 | 374 |

Next, we generated heatmaps to visualize ATAC-seq read coverage and CLAMP and ZLD ChIP-seq occupancy at their related classes of loci in MTD (wt), *clamp-i* and *zld-i* (*zld-*) embryos **(Figure 5-figure supplement 1A)**. As expected, MTD and *clamp-i* embryo heatmaps revealed that CLAMP-related DA regions (DA with CLAMP, DA without CLAMP) show a significant decrease in accessibility in embryos lacking CLAMP. Regions dependent on ZLD to open (DA with ZLD, DA without ZLD) also show a significant accessibility reduction in the absence of ZLD. Moreover, ChIP-seq read enrichment for protein binding in each RNAi or germline clone embryo class corresponds to our classification.

Interestingly, both CLAMP and ZLD protein binding to chromatin was reduced when the other TF was depleted, especially at regions where the bound protein is not required for chromatin accessibility (non-DA) **(Figure 5-figure supplement 1A)**. For example, ZLD binding was reduced upon *clamp* RNAi at ZLD non-DA w/ ZLD regions. We also found that CLAMP is enriched at these ZLD non-DA regions, supporting our hypothesis that CLAMP might facilitate ZLD occupancy at some of these loci.

To assess the relationship between CLAMP and ZLD in regulating chromatin accessibility at loci bound by both factors, we next identified the subset of genomic loci (n=525) that co-bound both CLAMP and ZLD and have open zygotic chromatin **(Figure 5A, Table2** and **Table 2 – Source Data 1).** Next, we divided these co-bound loci into four types: one percent (n=5) of these loci show reduced accessibility after reducing levels of either CLAMP or ZLD (Type I, **Figure 5A**); 23 and 123 loci are specifically dependent on either CLAMP or ZLD for their accessibility, respectively (Type II & Type III, **Figure 5A)**; the majority (374 out of 525) of CLAMP ZLD co-bound loci remain open when either protein is absent (Type IV, **Figure 5A**), suggesting that CLAMP and ZLD function redundantly at some of these loci or that other TFs regulate their accessibility.

**Figure 5.**
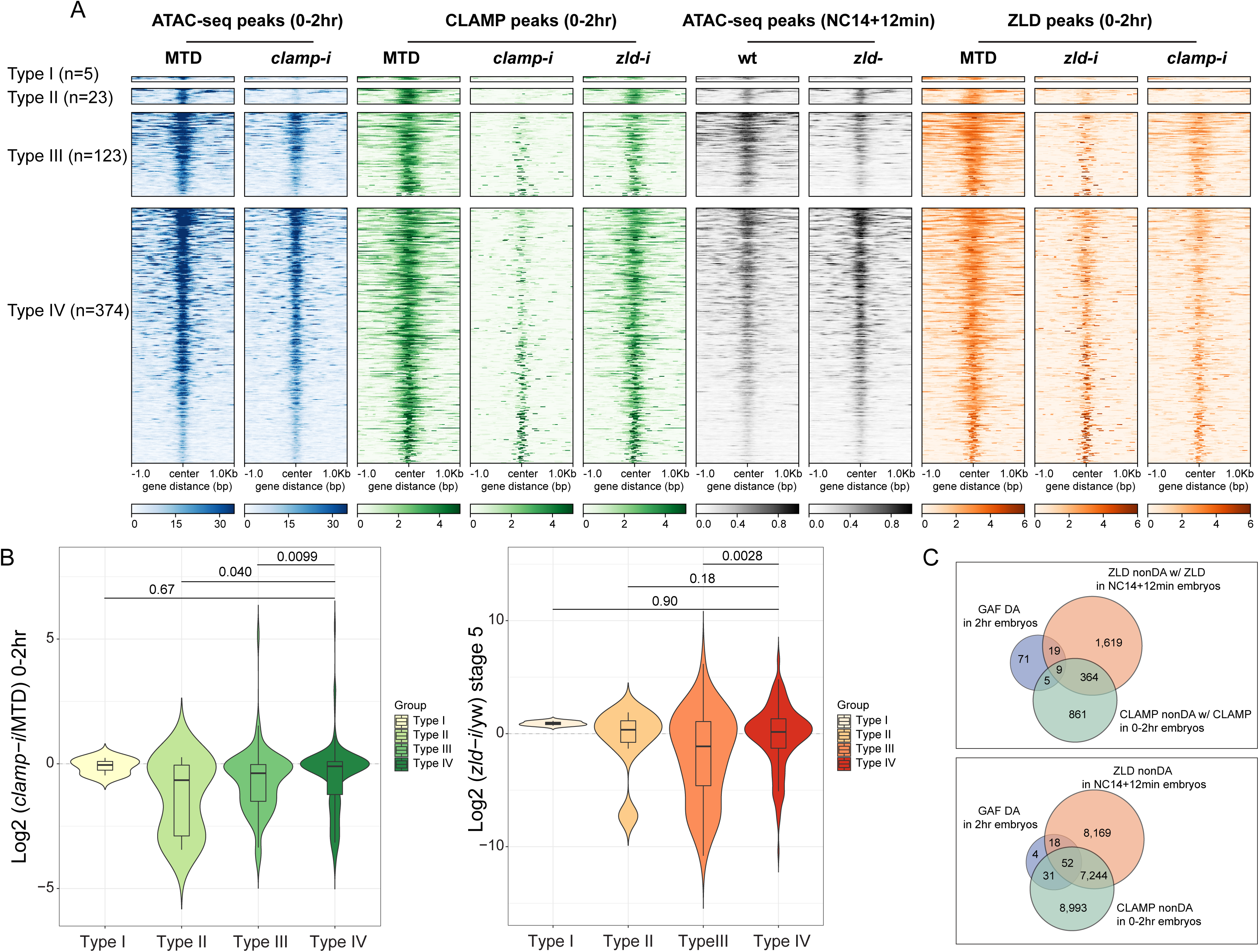
CLAMP and ZLD regulate gene expression via modulating chromatin accessibility. A. Four classes of CLAMP and ZLD co-bound peaks defined by combining ATAC-seq (this study or Hannon et al., 2017; Soluri et al., 2020) and ChIP-seq peaks in 0-2hr MTD and RNAi embryos. Data is displayed as a heatmap of z-score normalized ATAC-seq and ChIP-seq reads in a 2 kb region centered around each peak. Peaks in each class are arranged in order of decreasing *z*-scores in control MTD embryos.

▪ Type I (n=5): both DA, differentially accessible regions which depend on CLAMP or ZLD; has both proteins bound.
▪ Type II (n=23): CLAMP DA and ZLD non-DA, differentially accessible regions which depend on CLAMP, not on ZLD; has both proteins bound.
▪ Type III (n=123): ZLD DA and CLAMP non-DA, differentially accessible regions which depend on ZLD, not on CLAMP; has both proteins bound.
▪ Type IV (n=374): both non-DA, accessibility independent from CLAMP or ZLD but has both proteins bound. B. Left: Gene expression changes caused by maternal *clamp* RNAi (Rieder et al., 2017) in 0-2hr embryos at genes fall into four classes of CLAMP and ZLD co-bound peaks. *p*-values of significant expression changes among classes were calculated by Mann-Whitney U-test and noted on the plot. Right: Gene expression changes caused by maternal *zld* RNAi (Schulz et al., 2015) in stage5 embryos at genes fall into four classes of CLAMP and ZLD co-bound peaks. *p*-values of significant expression changes among classes were calculated by Mann-Whitney U-test and noted on the plot. C. Upper panel: Venn diagram showing the number of overlapping sites between GAF-dependent DA sites (Gaskill et al., 2021), ZLD non-DA with ZLD bound, and CLAMP non-DA with CLAMP bound peaks. Lower panel: Venn diagram showing the number of overlapping sites between GAF-dependent DA sites (Gaskill et al., 2021), ZLD non-DA, and CLAMP non-DA peaks.

Notably, at sites where CLAMP is required for chromatin accessibility (Type II, n=23), ZLD occupancy is entirely ablated in *clamp-i* embryos **(Figure 5A)**. CLAMP binding levels are also reduced after maternal *zld* RNAi at sites where ZLD is required for chromatin accessibility (Type III, n=123). Overall, we observed that CLAMP and/or ZLD occupancy is reduced at most of their co-bound regions when either one of the TFs is depleted, which is consistent with their inter-dependent binding relationship. Moreover, *clamp-i* has a stronger impact on ZLD binding than *zld-i* has on CLAMP binding.

To assess how CLAMP/ZLD-modulated chromatin accessibility impacts transcription, we examined the effect of maternal *clamp* (Rieder et al., 2017) or *zld* (Schulz et al., 2015) depletion on expression of genes that fall into the four types of CLAMP/ZLD co-occupied sites **(Figure 5B)**. We found that the expression levels of genes (Type II, n = 23) that require CLAMP for chromatin accessibility are significantly (*p* < 0.05, Mann-Whitney U-test) downregulated in embryos lacking CLAMP compared to the Type IV CLAMP and ZLD-independent group (n = 374) **(Figure 5B).** Genes (Type III, n = 123) dependent on ZLD for their accessibility also show a significant (*p* < 0.001, Mann-Whitney U-test) reduction in expression upon maternal CLAMP depletion, suggesting CLAMP also might contribute to the regulation of genes at which ZLD regulates chromatin accessibility, likely by increasing ZLD binding.

In embryos depleted for maternal ZLD (Schulz et al., 2015), we found genes that fall into the Type III ZLD-mediate chromatin accessibility group significantly (*p* < 0.001, Mann-Whitney U-test) decreased in expression **(Figure 5B),** compared with the Type IV (n = 374) group. Interestingly, genes within the CLAMP and ZLD-independent Type IV (n = 374) group do not show significant expression fold changes after depleting either maternal *clamp* or *zld*, indicating that CLAMP and ZLD function redundantly at these loci and/or other proteins play a major role in regulating chromatin accessibility and transcription of these genes.

Motif analysis demonstrates that CLAMP and ZLD motifs are enriched at genomic loci that are regulated by each factor as well as independent sites (Type IV), in addition to the motif for another GA-binding protein, GAF **(Figure 5-figure supplement 1B**). We next determined whether GAF alters chromatin accessibility at loci at which depletion of CLAMP or ZLD individually alters accessibility (Type IV) and are bound by all three factors. Indeed, we found that approximately 10% of loci that require GAF for their chromatin accessibility (n=104) (Gaskill et al., 2021) overlap with regions where depleting CLAMP or ZLD individually does not alter accessibility **(Figure 5C upper panel)**. When we do not require occupancy of ZLD and CLAMP at their non-DA sites, the overlap is approximately 97% with GAF-dependent loci **(Figure 5C lower panel)**. These results suggesting GAF might function at these CLMAP/ZLD independent sites, supporting a model in which multiple TFs coordinately regulate early zygotic chromatin accessibility during ZGA.

Together, our results reveal the CLAMP and ZLD regulate chromatin accessibility, which alters the occupancy of both factors and regulates zygotic transcription. Furthermore, GAF and/or other TFs might function at sites that are not altered by depleting CLAMP or ZLD individually, suggesting that multiple TFs promote chromatin accessibility during ZGA. It is also possible that CLAMP and ZLD are functionally redundant at the subset of genomic loci at which they regulate each other’s occupancy, but depleting either factor individually is not sufficient to alter chromatin and expression.

## DISCUSSION

Two questions central to early embryogenesis of all metazoans are how and where do early transcription factors work together to drive chromatin changes and zygotic genome activation. Here, we identified CLAMP as a pioneer-like transcription factor that is essential for early embryonic development, which directly binds to nucleosomal DNA **(Figure 1)**, establishes and/or maintains chromatin accessibility at promoters of genes that often encode other TFs **(Figure 2)**, facilitates the binding of ZLD to promoters **(Figure 3)** to regulate zygotic genome activation **(Figure 4** and **Figure 5)**. We further discovered that CLAMP and ZLD regulate each other’s binding during ZGA to increase binding to promoters of genes which often encode other early TFs. Also, we determined that both CLAMP and ZLD alter chromatin accessibility, which influences the binding of both TFs to their binding sites **(Figure 6)**. Overall, we provide new insight into how CLAMP and ZLD function together to enhance each other’s occupancy and increase chromatin accessibility, which drives zygotic genome activation.

**Figure 6.**
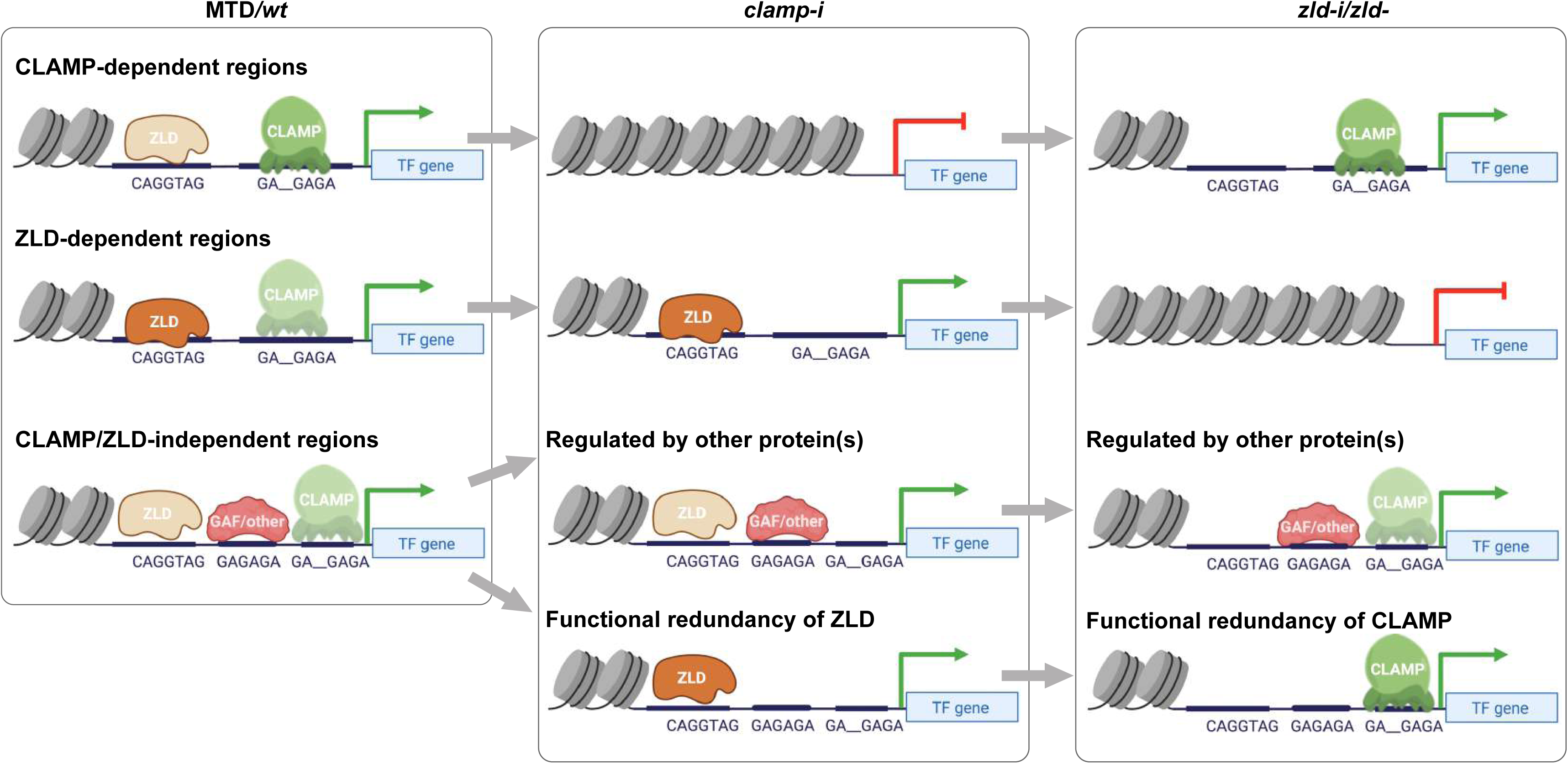
Model for how CLAMP and ZLD pioneer factor function together to define chromatin accessibility in early embryos. CLAMP and ZLD function together at promoters to regulate each other’s occupancy and gene expression of genes encoding other key TFs. We defined CLAMP and ZLD co-bound peaks in early embryos, which revealed roles for CLAMP and ZLD in defining chromatin accessibility and activating zygotic transcription at a subset of the zygotic genome. **CLAMP-dependent regions:** CLAMP promotes ZLD enrichment at these sites where CLAMP binding increases chromatin accessibility and regulates target gene expression. These sites are closed and lack binding of ZLD when maternal *clamp* is depleted, and they remain open and transcription is activated when maternal *zld* is depleted. **ZLD-dependent regions:** ZLD modulates chromatin opening and transcription at these sites that are bound by CLAMP but do not depend on CLAMP for chromatin accessibility. These sites are closed and lack binding of CLAMP when maternal *zld* is depleted, and they remain open and active when maternal *clamp* is depleted. **CLAMP/ZLD-independent regions:** GAF or other TFs open chromatin at locations co-bound by CLAMP and ZLD where chromatin accessibility is not altered when each factor is depleted individually. CLAMP and ZLD could also function redundantly at some of these loci. These sites remain accessible and transcriptionally active upon either maternal *zld* or *clamp* depletion.

### CLAMP and ZLD act together to define an open chromatin landscape and activate transcription in early embryos

We defined multiple classes of CLAMP-dependent and ZLD-dependent genomic loci in early embryos, which provides insight into how CLAMP and ZLD regulate chromatin accessibility and zygotic transcription during ZGA **(Figure 6)**: 1) CLAMP promotes ZLD enrichment at sites where CLAMP increases chromatin accessibility and further regulates ZLD target gene expression. These loci remain open and transcriptionally active even upon ZLD depletion. 2) ZLD facilitates CLAMP occupancy at sites where ZLD regulates chromatin accessibility and promotes CLAMP target gene expression. When maternal CLAMP is depleted, these loci remain accessible and genes are actively transcribed. 3) GAF and/or other TFs could play major roles in opening chromatin at locations co-bound by CLAMP and ZLD but that are not altered in accessibility after depleting CLAMP or ZLD individually. CLAMP and ZLD could also function redundantly at some of these loci because they alter each other’s occupancy at these loci but do not change accessibility or expression after depletion of either maternal CLAMP or ZLD individually. Overall, our data suggest that CLAMP functions with ZLD to regulate chromatin accessibility and gene expression of the early zygotic genome.

Although we have demonstrated an instrumental role for CLAMP in defining a subset of the open chromatin landscape in early embryos, our data show that CLAMP does not increase chromatin accessibility at promoters of all zygotic genes independent of ZLD. Consistent with our results in the early embryo, CLAMP regulates chromatin accessibility at only a few hundred genomic loci in male S2 (258 sites) and female Kc (102 sites) cell lines (Albig et al., 2019). Unlike ZLD, which plays a global role in regulating chromatin accessibility at promoters throughout the genome, depletion of CLAMP alone mainly drives changes at promoters of specific genes that often encode transcription factors that are important for early development, consistent with phenotypic data.

Moreover, ZLD binding and/or chromatin accessibility is not regulated by maternal depletion of CLAMP at all GA-rich sites in the genome. GAF is also enriched at these same ZLD-bound regions where ZLD is not required for chromatin accessibility (Schulz et al., 2015; Gaskill et al., 2021). Both CLAMP and GAF are deposited maternally (Rieder et al., 2017; Hamm et al., 2017) and bind to similar GA-rich motifs (Kaye et al., 2018). To test whether GAF compensates for the depletion of CLAMP or ZLD, we tried to perform GAF RNAi in the current study to prevent GAF from compensating for CLAMP depletion. However, we and other laboratories could not achieve depletion of GAF in early embryos by RNAi, likely due to autoregulation of its own promoter and its prion-like self-perpetuating function (Tariq et al., 2013).

We previously demonstrated that competition between CLAMP and GAF at GA-rich binding sites is essential for MSL complex recruitment in S2 cells (Kaye et al., 2018). Furthermore, CLAMP excludes GAF at the histone locus which co-regulates genes that encode the histone proteins (Rieder et al., 2017). However, we also observed synergistic binding between CLAMP and GAF at many additional binding sites (Kaye et al., 2018). The relationship between CLAMP and GAF in early embryos remains unclear. It is very possible that the competitive relationship has not been established in early embryos, since dosage compensation has not yet been initiated (Prayitno et al., 2019). Using GAF-dependent loci defined by Gaskill et al. (2021), we found that genomic loci where GAF functions largely overlap with regions where depletion of CLAMP or ZLD alone does not alter chromatin accessibility, indicating that GAF may function independently of CLAMP or ZLD or is functionally redundant. Future studies are required to distinguish between these models by examining how GAF and CLAMP affect each other’s binding to co-bound loci and simultaneously eliminating both factors.

The GA-rich sequences targeted by CLAMP and GAF are distinct from each other *in vivo* and *in vitro*. GAF binding sites typically have 3.5 GA repeats; however, GAF is able to bind to as few as three bases (GAG) within the *hsp70* promoter and *in vitro* (Wilkins and Lis, 1999). In contrast, CLAMP binding sites contain an 8-bp core with a less well-conserved second GA dinucleotide within the core (GA GAGA) (Alekseyenko et al., 2008). CLAMP binding sites also include a GAGAG pentamer at a lower frequency than GAF binding sites, and flanking bases surrounding the 8-bp core are critical for CLAMP binding (Kaye et al., 2018). Therefore, GAF and CLAMP may have overlapping and non-overlapping functions at different loci, tissues, or developmental stages. Moreover, another TF, Pipsqueak (Psq) also binds to sites containing the GAGAG motif, and has multiple functions during oogenesis andembryonic pattern formation and functions with Polycomb in three-dimensional genome organization (Lehmann et al., 1998; Gutierrez-Perez et al., 2019). In the future, an optogenetic inactivation approach could be used to remove CLAMP, GAF, and/or Psq simultaneously in a spatial and temporal manner (McDaniel et al., 2019).

### CLAMP and ZLD regulate each other’s binding via their own motifs

ZLD is an essential TF that regulates activation of the first set of zygotic genes during the minor wave of ZGA and thousands of genes transcribed during the major wave of ZGA at nuclear cycle 14 (Liang et al., 2008; Harrison et al., 2011). ZLD also establishes and maintains chromatin accessibility of specific regions and facilitates transcription factor binding and early gene expression (Sun et al., 2015; Schulz et al., 2015). CLAMP regulates histone gene expression (Rieder et al., 2017), X chromosome dosage compensation (Soruco et al., 2013), and establishes/maintains chromatin accessibility (J. Urban et al., 2017). Nonetheless, it remained unclear whether and how CLAMP and ZLD functionally interact during ZGA. Here, we demonstrate that CLAMP and ZLD function together at a subset of promoters that often encode other transcriptional regulators.

ZLD regulates CLAMP occupancy earlier than CLAMP regulates ZLD occupancy. Genomic loci at which CLAMP is dependent on ZLD early (0-2hr) in development often become independent from ZLD later (2-4hr), with the caveat that ZLD depletion is not as effective later in development. Therefore, CLAMP may require the pioneering activity of ZLD to access specific loci before ZGA, but ZLD may no longer be necessary once CLAMP binding is established. Also, our results suggest that CLAMP is a potent regulator of ZLD binding, especially in 2-4hr embryos. ZLD can bind to many more promoter regions at 0-2 hours, while CLAMP mainly binds to introns early in development but occupies promoters later at 2-4 hours. Therefore, CLAMP may require ZLD to increase chromatin accessibility of these promoter regions (Schulz et al., 2015).

In addition to its role in embryonic development, CLAMP also plays an essential role in targeting the MSL male dosage compensation complex to the X-chromosome (Soruco et al., 2013). *Drosophila* embryos initiate X-chromosome counting in nuclear cycle 12 and start the sex determination cascade prior to the major wave of ZGA at nuclear cycle 14 (Gergen, 1987; Bosch et al., 2006). However, most dosage compensation is initiated much later in embryonic development (Prayitno et al., 2019). Our data support a model in which CLAMP functions early in the embryo prior to MSL complex assembly to open up specific chromatin regions for MSL complex recruitment (J. Urban et al., 2017; Rieder et al., 2019). Moreover, ZLD likely functions primarily as an early pioneer factor, whereas CLAMP has pioneer-like functions in both early- and late-ZGA embryos. Consistent with this hypothesis, CLAMP binding is enriched at both early and late zygotic genes. In contrast, ZLD binding binds more frequently to early genes, suggesting a that there may be a sequential relationship between occupancy of these two TFs at some loci during early embryogenesis.

The different characteristics of dependent and independent CLAMP and ZLD binding sites also provide insight into how early transcription factors work together to regulate ZGA. At dependent sites, there are often relatively broad peaks of CLAMP and ZLD that are significantly enriched for clusters of motifs for the required protein. Our CLAMP gel shift assays and those previously reported (Kaye et al., 2018) also show multiple shifted bands consistent with possible multimerization. CLAMP contains two central disordered prion-like glutamine-rich regions (Kaye et al., 2018), a domain that is critical for transcriptional activation and multimerization *in vivo* in several TFs, including GAF (Wilkins and Lis, 1999). Moreover, glutamine-rich repeats alone can be sufficient to mediate stable protein multimerization *in vitro* (Stott et al., 1995). Therefore, it is reasonable to hypothesize that the CLAMP glutamine-rich domain also functions in CLAMP multimerization.

In contrast, ZLD fails to form dimers or multimers (Hamm et al., 2015, 2017), indicating that ZLD most likely binds as a monomer. There is no evidence that CLAMP and ZLD have any direct protein-protein interaction at sites where they depend on each other to bind. For example, mass spectrometry results that identified dozens of CLAMP-associated proteins did not identify ZLD (J. A. Urban et al., 2017). No data has validated any protein-protein interactions of ZLD with itself as a multimer or between ZLD and any other TFs (Hamm et al., 2017). In the future, simultaneous ablation of maternal CLAMP and ZLD will allow analysis of potential functional redundancy at a subgroup of genomic loci. Our study suggests that regulating the chromatin landscape in early embryos to drive ZGA requires the function of multiple pioneer transcription factors.

## MATERIALS AND METHODS

### Recombinant Protein Expression and Purification of CLAMP

MBP-tagged CLAMP DBD was expressed and purified as described previously (Kaye et al., 2018). MBP-tagged (pTHMT, Peti and Page, 2007) full-length CLAMP protein was expressed in *Escherichia coli* BL21 Star (DE3) cells (Life Technologies). Bacterial cultures were grown to an optical density of 0.7 to 0.9 before induction with 1 mM isopropyl-β-D-1-thiogalactopyranoside (IPTG) for 4 hrs at 37°C.

Cell pellets were harvested by centrifugation and stored at −80°C. Cell pellets were resuspended in 20 mM Tris, 1M NaCl, 0.1mM ZnCl2, 10 mM imidazole pH 8.0 with one EDTA-free protease inhibitor tablet (Roche) and lysed using an Emulsiflex C3 (Avestin). The lysate was cleared by centrifugation at 20,000 rpm for 50 min at 4°C, filtered using a 0.2 μm syringe filter, and loaded onto a HisTrap HP 5 mL column. The protein was eluted with a gradient from 10 to 300 mM imidazole in 20 mM Tris, 1.0 M NaCl pH 8.0, and 0.1mM ZnCl2. Fractions containing MBP-CLAMP full-length were loaded onto a HiLoad 26/600 Superdex 200 pg column equilibrated in 20 mM Tris 1.0 M NaCl pH 8.0. Fractions containing full-length CLAMP were identified by SDS-PAGE and concentrated using a centrifugation filter with a 10 kDa cutoff (Amicon, Millipore) and frozen as aliquots.

### *In vitro* assembly of nucleosomes

The 240 bp 5C2 DNA fragment used for nucleosome *in vitro* assembly was amplified from 276 bp 5C2 fragments (50ng/ul, IDT gBlocks Gene Fragments) by PCR (see 276 bp 5C2 and primer sequences below) using OneTaq Hot Start 2X Master Mix (New England Biolabs). The DNA was purified using the PCR clean-up kit (Qiagen) and concentrated to 1ug/ul by SpeedVac Vacuum (Eppendorf). The nucleosomes were assembled using the EpiMark® Nucleosome Assembly Kit (New England Biolabs) following the kit’s protocol.

5C2 (276 bp), **bold** sequences are CLAMP-binding motifs, underlined sequences are primer binding sequences: TCGACGACTAGTTTAAAGTTA<u>TTGTAGTTCTTAGAGCAGAATGT</u>ATTTTAAATATCAATGTTT CGATGTAGAAATTGAATGGTTTAAATCACGTTCACACAACTTA**GAAAGAGATAG**CGATGGC GGTGT**GAAAGAGAGCGAGATAG**TTGGAAGCTTCATG**GAAATGAAAGAGAGGTAG**TTTTT GGAAATGAAAGTTGTACTAGAAATAAGTATTTTATGTATATAGAATATCGAAGTACAG<u>AAATT CGAAGCGATCTCAAC</u>TTGAATATTATATCG

**Primers for 5C2 region (product is 240bp):**

Forward: TTGTAGTTCTTAGAGCAGAATGT

Reverse: GTTGAGATCGCTTCGAATTT

### Electromobility shift assays

DNA or nucleosome probes at 35nM (700fmol/reaction) were incubated with MBP-tagged CLAMP DBD protein or MBP-tagged full-length CLAMP protein in a binding buffer. The binding reaction buffer conditions are similar to conditions previously used to test ZLD nucleosome binding (McDaniel et al. 2019) in 20 ul total volume: 7.5ul BSA/HEGK buffer (12.5 mM HEPES, PH 7.0, 0.5 mM EDTA, 0.5 mM EGTA, 5% glycerol, 50 mM KCl, 0.05 mg/ml BSA, 0.2 mM PMSF, 1 mM DTT, 0.25 mM ZnCl2, 0.006% NP-40,) 10 ul probe mix (5 ng poly[d-(IC)], 5 mM MgCl2, 700 fmol probe), and 2.5 ul protein dilution (0.5uM, 1uM, 2.5uM) at room temperature for 60 min. Reactions were loaded onto 6% DNA retardation gels (ThermoFisher) and run in 0.5X Tris–borate–EDTA buffer for 2 hours. Gels were post stained with GelRed Nucleic Acid Stain (Thermo Scientific) for 30 min and visualized using the ChemiDoc MP imaging system (BioRad).

### Fly stocks and crosses

To deplete maternally deposited *clamp* or *zld* mRNA throughout oogenesis, we crossed a maternal triple driver (MTD-GAL4, Bloomington, #31777) line (Ni et al., 2011) with a Transgenic RNAi Project (TRiP) *clamp* RNAi line (Bloomington, #57008), a TRiP *zld* RNAi line (from C. Rushlow lab) or *egfp* RNAi line (Bloomington, #41552). The *egfp* RNAi line was used as control in smFISH immunostaining and imaging experiments. The MTD-GAL4 line alone was used as the control line in ATAC-seq and ChIP-seq experiments.

Briefly, the MTD-GAL4 virgin females (5-7 day old) were mated with TRiP UAS-RNAi males to obtain MTD-Gal4/UAS-RNAi line daughters. The MTD drives RNAi during oogenesis in these daughters. Therefore, the targeted mRNA is depleted in their eggs. Then MTD-Gal4/UAS-RNAi daughters were mated with males to produce embryos with depleted maternal *clamp* or *zld* mRNA and used for ATAC-seq and ChIP-seq experiments. The embryonic phenotypes of the maternal *zld*^-^ TRiP RNAi line were confirmed previously (Sun et al., 2015). Maternal *clamp*^-^ embryonic phenotypes of the TRiP *clamp* RNAi line were confirmed by immunofluorescent staining in our study. Moreover, we validated CLAMP or ZLD protein knockdown in early embryos by western blotting using the Western Breeze kit (Invitrogen) and measured *clamp* and *zld* mRNA levels by qRT-PCR (**Figure 1 – figure supplement 1B, C** and **Figure 1 Source Data 1-2**).

### Embryo collections

To optimize egg collections, young (5-7 day old) females and males were mated. To ensure mothers do not lay older embryos during collections, we first starved flies for 2 hours in the empty cages and discarded the first 2-hour grape agar plates with yeast paste (Plate set #0). When we collected eggs for the experiments, we put flies in the cages with grape agar plates (Plate set #1) with yeast paste for egg laying for 2 hours. Then, we replaced Plate set #1 with a new set of plates (Plate set #2) at the 2hr time point. We kept Plate set #1 embryos (without any adult flies) to further develop for another 2 hours to obtain 2-4hr embryos. At the same time, we obtained newly laid 0-2hr embryos from Plate set #2. Therefore, this strategy successfully prevented cross contamination between 0-2hr embryos (Plate set #2) and the 2-4hr embryos (Plate set #1).

### smFISH, Immunostaining and Imaging

For whole embryo single molecule fluorescence *in situ* hybridization (smFISH) and immunostaining and subsequent imaging, standard protocols were used (Little and Gregor, 2018). smFISH probes complementary to *run* were a gift from Thomas Gregor, and those complementary to *eve* were a gift from Shawn Little. The concentrations of the different dyes and antibodies were as follows: Hoechst (Invitrogen, 3µg/ml), Anti-NRT (Developmental Studies Hybridoma Bank BP106, 1:10), AlexaFluor secondary antibodies (Invitrogen Molecular Probes, 1:1000). Imaging was done using a Nikon A1 point-scanning confocal microscope with a 40X oil objective. Image processing and intensity measurements were done using Image J software (NIH). Figures were assembled using Adobe Photoshop CS4.

### ATAC-seq in embryos

We conducted ATAC-seq following the protocol from Blythe and Wieschaus, (2016). 0-2hr or 2-4hr embryos were laid on grape agar plates, dechorionated by 1 min exposure to 6% bleach (Clorox) and then washed 3 times in deionized water. We homogenized 10 embryos and lysed them in 50 ul lysis buffer (10mM Tris 7.5, 10mM NaCl, 3mM MgCl2, 0.1% NP-40). We collected nuclei by centrifuging at 500 g at 4°C and resuspended nuclei in 5 ul TD buffer with 2.5 ul Tn5 enzyme (Illumina Tagment DNA TDE1 Enzyme and Buffer Kits). We incubated samples at 37°C for 30 min at 800 rpm (Eppendorf Thermomixer) for fragmentation, and then purified samples with Qiagen MinElute columns before PCR amplification. We amplified libraries by adding 10 ul DNA to 25 ul NEBNext HiFi 2x PCR mix (New England Biolabs) and 2.5 ul of a 25 uM solution of each of the Ad1 and Ad2 primers. We used 13 PCR cycles to amplify samples from 0-2hr embryos and 12 PCR cycles to amplify samples from 2-4hr embryos. Next, we purified libraries with 1.2x Ampure SPRI beads. We performed three biological replicates for each genotype (n=2) and time point (n=2). We measured the concentrations of 12 ATAC-seq libraries by Qubit and determined library quality by Bioanalyzer. We sequenced libraries on an Illumina Hi-seq 4000 sequencer at GeneWiz (South Plainfield, NJ) using the 2×150-bp mode. ATAC-seq data is deposited at NCBI GEO and the accession number is GSE152596.

### Chromatin Immunoprecipitation-sequencing (ChIP-seq) in embryos

We performed ChIP-seq as previously described Blythe and Wieschaus, (2015). We collected and fixed ∼100 embryos from each MTD-GAL4 and RNAi cross 0-2hr or 2-4hr after egg lay. We used 3 ul of rabbit anti-CLAMP (Soruco et al., 2013) and 2 ul rat anti-ZLD (from C. Rushlow lab) per sample. We performed three biological ChIP replicates for each protein (n=2), genotype (n=3) and time point (n=2). In total, we prepared 36 libraries using the NEBNext Ultra ChIP-seq kit (New England Biolabs) and sequenced libraries on the Illumina HiSeq 2500 sequencer using the 2×150-bp mode. ChIP-seq data is deposited at NCBI GEO and the accession number is GSE152598.

### Computational analyses

#### ATAC-seq analysis

Prior to sequencing, the Fragment Analyzer showed the library top peaks were in the 180-190bp range, which is comparable to the previously established embryo ATAC-seq protocol (Haines, 2017). Demultiplexed reads were trimmed of adapters using TrimGalore (Krueger, 2017) and mapped to the *Drosophila* genome dm6 version using Bowtie2 (v. 2.3.0) with option --very-sensitive, --no-mixed, --no-discordant, --dovetail -X 2000 -k 2. We used Picard tools (v. 2.9.2) and SAMtools (v.1.9, Li et al., 2009) to remove the reads that were unmapped, failed primary alignment, or duplicated (-F 1804), and retain properly paired reads (-f 2) with MAPQ >30. After quality trimming and mapping, the Picard tool reported the mean fragment sizes for all ATAC-seq mapped reads is between 125-161bp. As expected, we observed three classes of peaks: 1) A sharp peak at <100 bp (open chromatin); 2) A peak at ∼200bp (mono-nucleosome); 3) Other larger peaks (multi-nucleosomes).

After mapping, we used Samtools to select a fragment size <= 100bp within the bam files to focus on open chromatin. Peak regions for open chromatin regions were called using MACS2 (version 2.1.1, Zhang et al., 2008) with parameters -f BAMPE -g dm --call-summits. ENCODE blacklist was used to filter out problematic regions in dm6 (Amemiya et al., 2019). Bam files and peak bed files were used in DiffBind v.3.12 (Stark and Brown, 2019) for count reads (dba.count), library size normalization (dba.normalize) and calling (dba.contrast) differentially accessible (DA) region with the DESeq2 method. Peak regions (201bp) were centered by peak summits and extended 100bp on each side. Sites were defined as DA with statistically significant differences between conditions using absolute cutoffs of FC > 0.5 and FDR < 0.1. We report all accessible peaks from DiffBind in **Figure 2-supplementary table 1.**

We used DeepTools (version 3.1.0, Ramírez et al., 2014) to generate enrichment heatmaps (CPM normalization), and average profiles were generated in DeepStats (Gautier RICHARD, 2020). We used 1x depth (reads per genome coverage, RPGC) normalization in Deeptools bamCoverage for making the coverage Bigwig files and uploaded to IGV (Robinson et al., 2011) for genomic track visualizations. Homer (v 4.11, Givler and Lilienthal, 2005) was used for *de novo* motif searches. Visualizations and statistical tests were conducted in R (R Core Team, 2014). Specifically, we annotated peaks to their genomic regions using R packages Chipseeker (Yu et al., 2015) and we performed gene ontology enrichment analysis using clusterProfiler (Yu et al., 2012). Boxplot and violin plots were generated using ggplot2 (Wickham, 2009) package.

#### ChIP-seq analysis

Briefly, we trimmed ChIP sequencing raw reads with TrimGalore (Krueger, 2017) with a minimal phred score of 20, 36 bp minimal read length, and Illumina adaptor removal. We then mapped cleaned reads to the *D. melanogaster* genome (UCSC dm6) with Bowtie2 (v. 2.3.0) with the –very-sensitive-local flag feature. We used Picard tools (v. 2.9.2) and SAMtools (v.1.9, Li et al., 2009) to remove the PCR duplicates. We used MACS2 (version 2.1.1, Zhang et al., 2008) to identify peaks with default parameters and MSPC (v.4.0.0, Jalili et al., 2015) to obtain consensus peaks from 3 replicates. The peak number for each sample was summarized in Table 4. ENCODE blacklist was used to filter out problematic regions in dm6 (Amemiya et al., 2019). We identified differential binding (DB) and non-differential binding (non-DB) between MTD and RNAi samples using DiffBind (v. 3.10, Stark and Brown, 2019) with the DESeq2 method. Peak regions (501bp) were centered by peak summits and extended 250bp on each side. The DB and non-DB peak numbers are summarized in **Table 1**. Differential binding was defined with absolute FC > 0.5 and FDR < 0.05 (**Table 1 – Source Data 1**).

We used DeepTools (version 3.1.0, Ramírez et al., 2014) to generate enrichment heatmaps and average profiles. Bigwig files were generated with DeepTools bamCompare (scale factor method: SES; Normalization: log2) and uploaded to IGV (Robinson et al., 2011) for genomic track visualization. We used Homer (v 4.11, Givler and Lilienthal, 2005) for *de novo* motif searches and genomic annotation. Intervene (Khan and Mathelier, 2017) was used for intersection and visualization of multiple peak region sets. Visualizations and statistical tests were conducted in R (R Core Team, 2014). Specifically, we annotated peaks to their genomic regions using the R package Chipseeker (Yu et al., 2015) and we did gene ontology enrichment analysis using clusterProfiler (Yu et al., 2012). Boxplots and violin plots were generated using the ggplot2 (Wickham, 2009) package.

#### ATAC-seq and ChIP-seq data integration

We used Bedtools (Quinlan and Hall, 2010) intersection tool to intersect peaks in CLAMP ChIP-seq binding regions with CLAMP DA or non-DA peaks. Based on the intersection of the peaks, we defined 4 types of CLAMP related peaks: 1) DA with CLAMP, 2) DA without CLAMP, 3) Non-DA with CLAMP, 4) Non-DA without CLAMP. Similarly, we defined ZLD related peaks by intersecting ZLD DA or non-DA peaks and ATAC-seq datasets (Hannon et al., 2017; Soluri et al., 2020) from wildtype (wt) and *zld* germline clone (*zld-*) embryos at the nuclear cycle14 +12 min stage. Specifically, we defined four classes of genomic loci for ZLD-related classes: 1) DA with ZLD, 2) DA without ZLD, 3) Non-DA with ZLD, 4) Non-DA, without ZLD. We used DeepTools (version 3.1.0, Ramírez et al., 2014) to generate enrichment heatmaps for each subclass of peaks. Peaks locations in each CLAMP or ZLD-related category were summarized in **Table 2 - Source Data 1**.

#### ATAC-seq and RNA-seq data integration

We annotated genes near differential (down-DA) ATAC-seq peaks in R using detailRanges function from the csaw package (Lun and Smyth, 2016). Then we plotted expression of genes using previously published RNA-seq data (Rieder et al., 2017).

#### ChIP-seq and RNA-seq data integration

To define strong, weak, and unbound genes close to peaks in CLAMP or ZLD ChIP-seq data, we used the peak binding score reported in MACS2 −log10(p-value) of 100 as a cut off value. We defined the following categories: 1) Strong binding peaks: score greater than 100; 2) Weak binding peak: score lesser than 100; 3) Unbound peaks: the rest of the peaks that are neither strong or weak. Then, we annotated all peaks using Homer annotatePeaks (v 4.11, Givler and Lilienthal, 2005). We then obtained the log2 fold change (*clamp-i/*MTD or *zld-i/yw*) of gene expression in the RNA-seq dataset for each protein binding group: CLAMP (Rieder et al., 2017) or ZLD (Schulz et al., 2015). Boxplots and violin plots were generated using the ggplot2 (Wickham, 2009) package.

#### Datasets

RNA-seq datasets from wild type and maternal *clamp* depletion by RNAi were from GSE102922 (Rieder et al., 2017). RNA-seq datasets from *yw* wild type and *zld* maternal RNAi were from GSE65837 (Schulz et al., 2015). ATAC-seq data from wild type and *zld* germline clones were from GSE86966 (Hannon et al., 2017). Processed ATAC-seq data identifying differential peaks between wild type and *zld* germline mutations were from Soluri et al., (2020).

#### Data accessibility for reviewers only

ATAC-seq (GSE152596) and ChIP-seq (GSE152598) are under GEO SuperSeries GSE152613. To review GEO accession GSE152613:

Go to https://www.ncbi.nlm.nih.gov/geo/query/acc.cgi?acc=GSE152613 Enter token ihevsmiqnxexrod into the box

## ACKNOWLEDGEMENTS

We thank Melissa Harrison, Tyler Gibson and Marissa Gaskill for sending the GAF-dependent region bed file and helpful discussions. This work was supported by NIH grant F32GM109663, K99HD092625 and R00HD092625 to Dr. Leila Rieder and R35GM126994 to Dr. Erica Larschan, and in part by NSF grant 1845734 and NIH grant R01GM118530 to Dr. Nicolas L. Fawzi.

## AUTHOR CONTRIBUTIONS

Conceptualization, L.E.R., J.E.D. and E.N.L.; Methodology, J.E.D., L.E.R. and E.N.L; ChIP-seq Experiment, L.E.R.; ATAC-seq Experiment, J.E.D.; Initial Analysis, W.J.III; Formal Analysis, J.E.D.; Microscopy, immunostaining and smFISH: M.M.C. and G.D. Protein Expression: M.M., S.W., and N.L.F.; Gel-shift Experiment, A.H and J.E.D.; Investigation, J.E.D; Data Curation, J.E.D; Writing--Original Draft, J.E.D. and E.N.L.; Writing--Review & Editing, J.E.D., L.E.R. and E.N.L.; Visualization, J.E.D.; Funding Acquisition, L.E.R., E.N.L. and N.L.F.

## DECLARATION OF INTERESTS

The authors declare no competing interests.

## Supplemental Figure Legends

**Figure 1-figure supplement 1.**
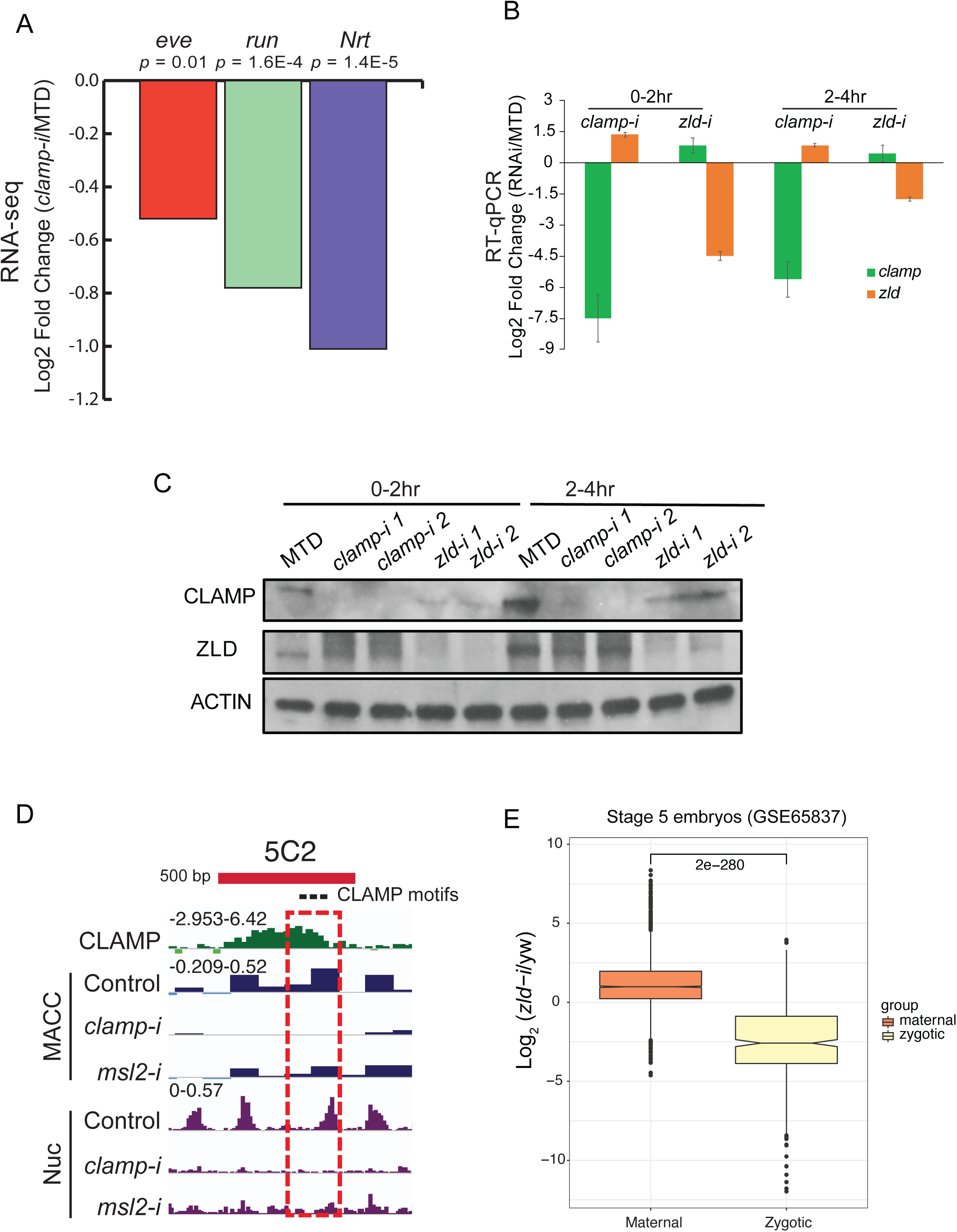
Novel pioneer factor CLAMP is essential for early embryonic development. A. Expression of early zygotic genes (*even-skipped*, *runt,* and *neurotactin*) in MTD and *clamp-i* embryos measured by RNA-seq (Rieder et al., 2017). Log_2_ Fold Change and *p-value* were calculated using DESeq2. B. Expression of mRNAs (*clamp* and *zld*) in MTD, *clamp-i* and *zld-i* embryos in 0-2hr and 2-4hr embryos measured by RT-qPCR. Log_2_ fold change was calculated using the ΔΔCt method (Rao et al., 2013) and normalized to reference gene *pka*. C. Western blot of CLAMP, ZLD and reference control Actin in MTD, *clamp-i* and *zld-i* embryos in 0-2hr and 2-4hr embryos. MTD: MTD-Gal4 line. *clamp-i:* MTD-Gal4-*clamp* mRNAi line, *zld-i:* MTD-Gal4-*zld* mRNAi line. D. Genome browser tracks for a region of the CES 5C2 locus (red bar is 500bp) used to make *in vitro* reconstituted nucleosomes (Urban et al., 2017). Locations of three CLAMP binding motifs are marked as black dots. CLAMP ChIP-seq normalized sequencing reads are shown in green. MNase-seq MACC scores (dark blue) show chromatin accessibility in S2 cells in control (*egfp-i*), *clamp*, and *msl2* RNAi treatment. The nucleosome profile (Nuc) is shown in purple. The dashed red rectangle highlights the genomic region (240bp) used to reconstitute nucleosomes *in vitro*. E. Effect of maternal *zld* RNAi (Schulz et al., 2015) on maternally-deposited (orange) or zygotically-transcribed (yellow) gene expression log2 (*clamp*-i/MTD) in 0-2hr (left) or 2-4hr (right). Maternal vs. zygotic gene categories were as defined in Lott et al. (2011). *p*-values of significant expression changes between maternal and zygotic genes were calculated by Mann-Whitney U-test and noted on the plot.

**Figure 2-figure supplement 1.**
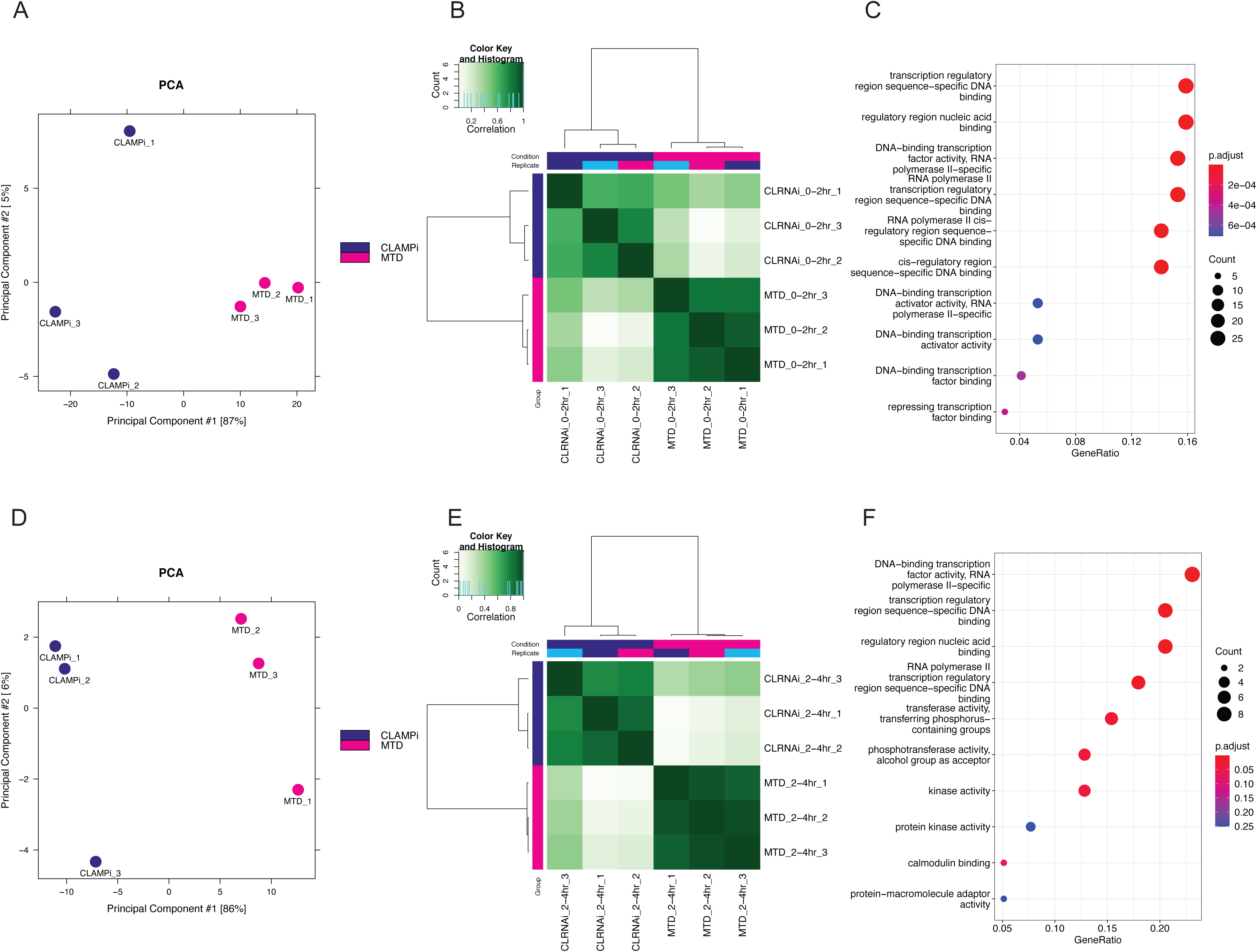
CLAMP regulates chromatin accessibility throughout ZGA. A. PCA plot for sample replicates in MTD and *clamp-i* embryos at 0-2hr time point. B. Pearson correlation heatmap of peaks dependent on CLAMP in MTD vs. *clamp-i* embryos at 0-2hr time point. C. GO terms for genes that require CLAMP for chromatin accessibility at 0-2hr time point. D. PCA plots for sample replicates in MTD and *clamp-i* embryos at 2-4hr time point. E. Pearson correlation heatmap of peaks dependent on CLAMP in MTD vs. *clamp-i* embryos at 2-4hr time point. F. GO terms for genes that require CLAMP for chromatin accessibility at 2-4hr time point.

**Figure 3-figure supplement 1.**
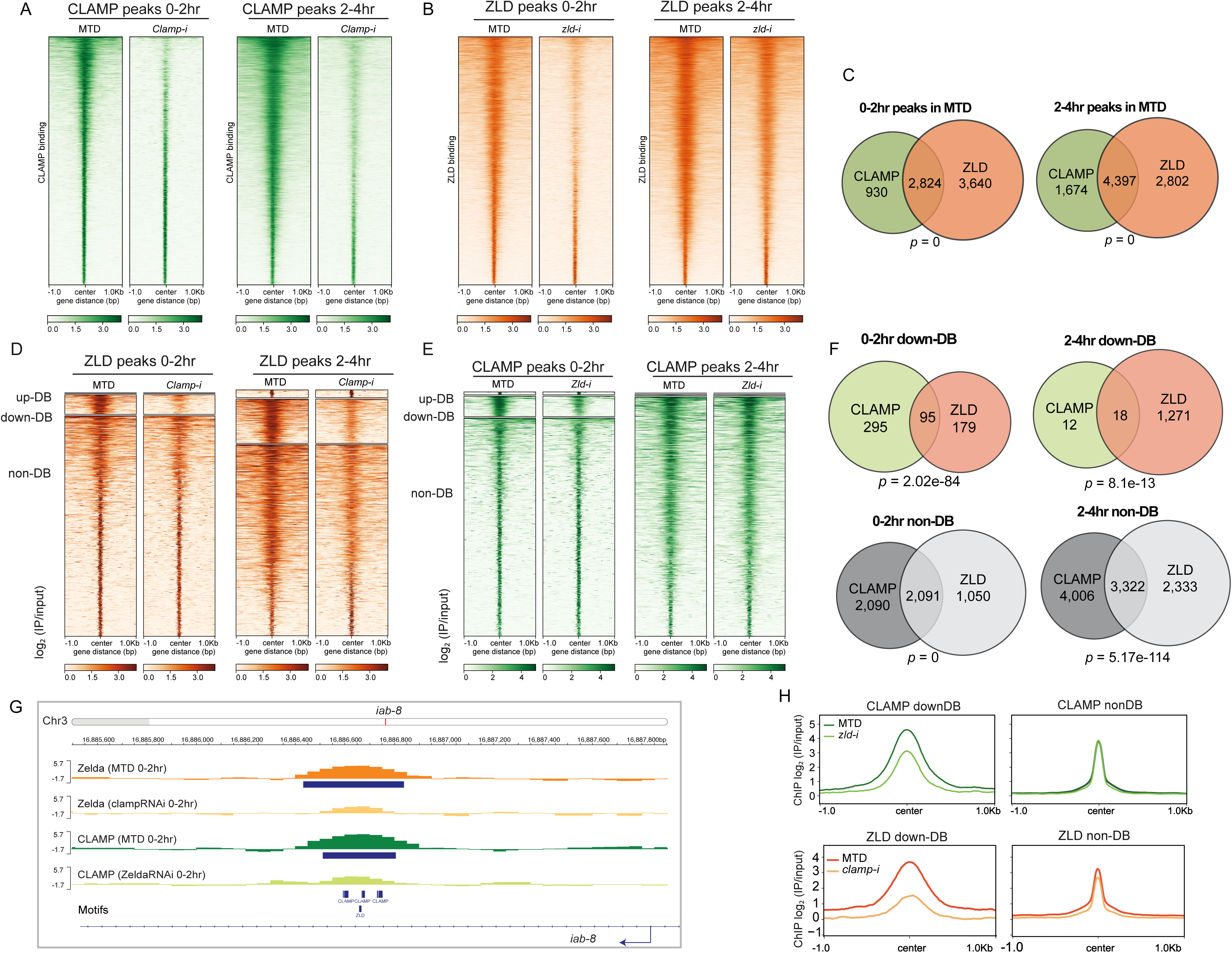
CLAMP and ZLD depend on each other for chromatin binding. A. CLAMP peaks in 0-2hr and 2-4hr MTD and maternal *clamp* RNAi embryos. Data is displayed as a heatmap of z-score normalized ChIP-seq (log2 IP/input) reads, in a 2 Kb region centered around each peak called in control MTD embryos. Peaks in each class are arranged in order of decreasing z-scores in control MTD embryos. B. ZLD peaks in 0-2hr and 2-4hr MTD and maternal *zld* RNAi embryos. Data is displayed as a heatmap of z-score normalized ChIP-seq (log2 IP/input) reads, in a 2 Kb region centered around each peak called in control MTD embryos. Peaks in each class are arranged in order of decreasing z-scores in control MTD embryos. C. CLAMP (green) and ZLD (orange) peaks and shared peaks where both CLAMP and ZLD are present in 0-2hr and 2-4hr embryos. *p*-values represent the significance (hypergeometric test, N= 15,682 total fly genes) of overlap. D. ZLD up-DB, down-DB and non-DB peaks in 0-2hr and 2-4hr MTD and maternal *clamp* RNAi embryos. Data is displayed as a heatmap of *z*-score normalized ChIP-seq (log2 IP/input) reads in a 2 Kb region centered around each peak. Peaks in each class are arranged in order of decreasing *z*-scores in control MTD embryos. E. CLAMP up-DB, down-DB and non-DB peaks in 0-2hr and 2-4hr MTD and maternal *zld* RNAi embryos. Data is displayed as a heatmap of z-score normalized ChIP-seq (log2 IP/input) reads in a 2 kb region centered around each peak. Peaks in each class are arranged in order of decreasing z-scores in control MTD embryos. F. Venn diagram showing the number of overlapping sites between ZLD and CLAMP down-DB or ZLD and CLAMP non-DB sites in 0-2hr and 2-4hr. *p*-values represent the significance (hypergeometric test, N= 15,682 total fly genes) of overlap. G. Example IGV views of genomic loci in *iab-8,* which CLAMP and ZLD were both down-DBs and dependent on each other to bind. H. Average profiles of ChIP-seq signal in log2 (IP/input) show the size of down-DB or non-DB peaks of CLAMP in MTD versus *zld*-i embryos (upper panel) and ZLD in MTD versus *clamp*-i embryos (lower panel).

**Figure 3-figure supplement 2.**
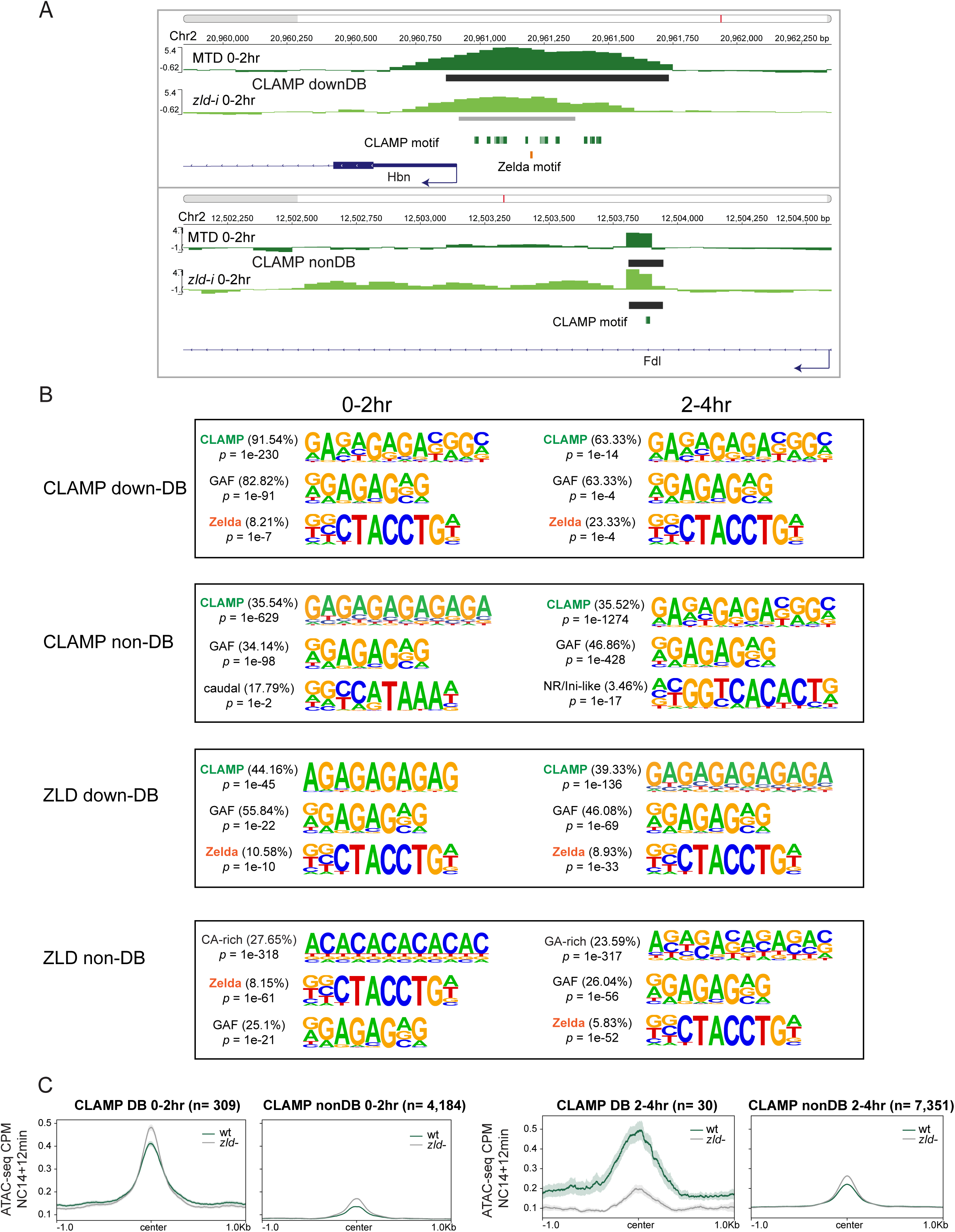
CLAMP and ZLD depend on each other for chromatin binding. A. Example IGV views of genomic loci: ChIP-seq in MTD and *zld*-i embryos at *Hbn* promoter region (upper panel) with CLAMP differential binding (down-DB) peak; *Fdl* (lower panel) with CLAMP non-differential binding (non-DB) peak at an intron. Peaks called by MACS2 are marked in dark grey (non-DB) and grey (down-DB). CLAMP and ZLD motifs are marked in green and orange respectively. B. Top three motifs called by Homer for CLAMP down-DB, CLAMP non-DB, ZLD down-DB, and ZLD non-DB peaks sites in 0-2hr and 2-4hr embryos. C. Average profiles of ATAC-seq signal coverage in NC14+12min wt and *zld*-embryos show chromatin accessibility at CLAMP down-DB (green line) and non-DB (grey line) sites defined in 0-2hr (left panel) or 2-4hr (right panel) MTD and *zld*-i embryos. Number of peaks is noted on the plot.

**Figure 5-figure supplement 1.**
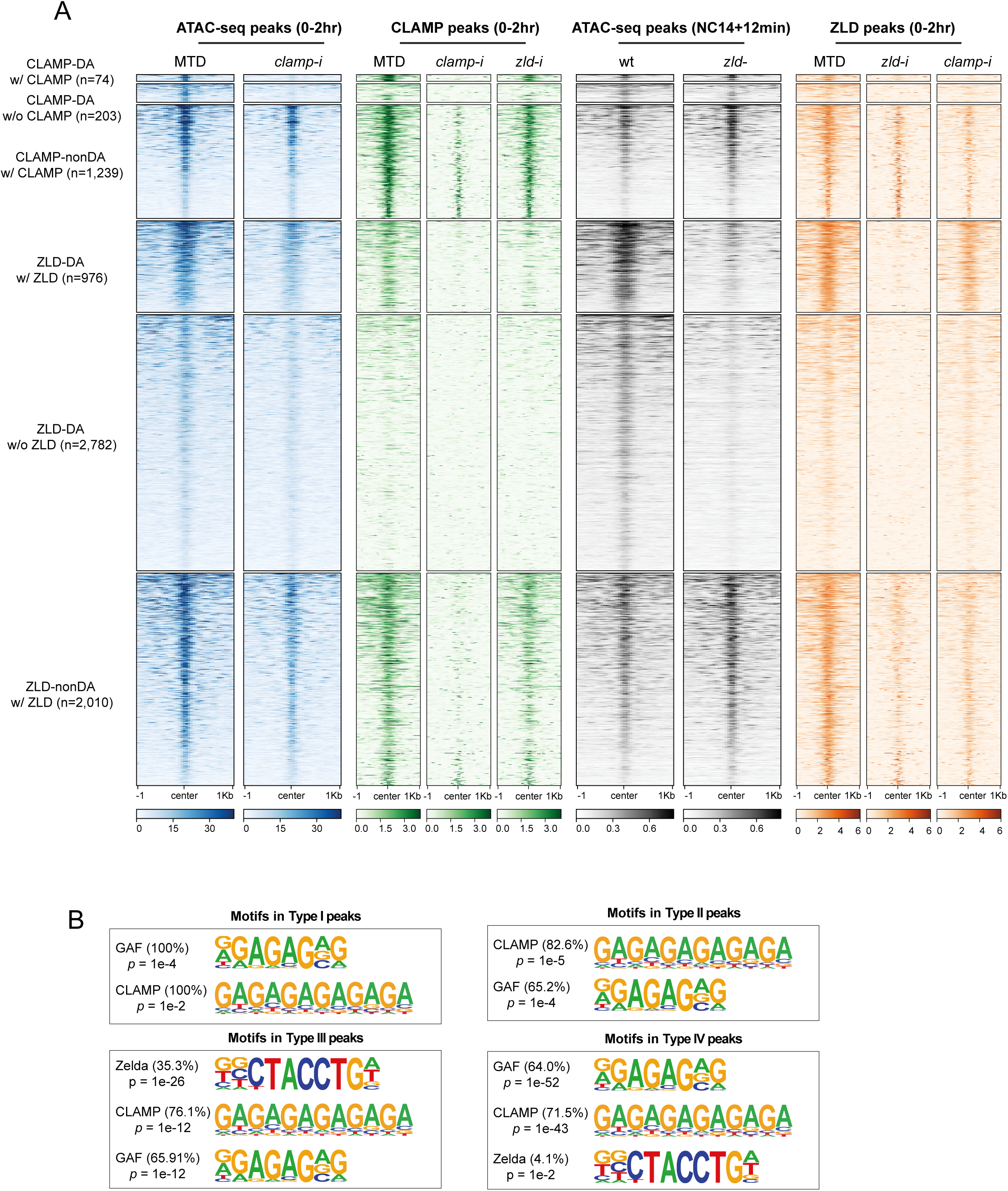
CLAMP-mediated chromatin accessibility is correlated with CLAMP and ZLD binding. A. CLAMP-related and ZLD-related peaks defined by combining ATAC-seq (this study or Hannon et al., 2017; Soluri et al., 2020) and ChIP-seq peaks in 0-2hr MTD and RNAi embryos. Data is displayed as a heatmap of z-score normalized ATAC-seq and ChIP-seq (log2 IP/input) reads, in a 2 Kb region centered around each peak. Peaks in each class are arranged in order of decreasing z-scores in control MTD embryos. Peak types and numbers are marked on plot:

▪ DA w/ CLAMP (n=74): differentially accessible regions that depend on CLAMP and has CLAMP binding.
▪ DA w/o CLAMP (n=203): differentially accessible regions that depend on CLAMP and has no CLAMP binding.
▪ Non-DA w/ CLAMP (n=1,238): not differentially accessible regions that have CLAMP binding.
▪ DA w/ ZLD (n=986): differentially accessible regions that depends on ZLD and has ZLD binding.
▪ DA w/o ZLD (n=2,797): differentially accessible regions that depends on ZLD and has no ZLD binding.
▪ NonDA w/ ZLD (n=2,301): non-differentially accessible regions which have ZLD binding. Note that Non-DA w/o CLAMP (n=15,081) and Non-DA w/o ZLD(n=17,758) peaks are omitted from this plot. B. Top motifs enriched in each type of CLAMP and ZLD co-bound regions. Enrichment *p*-value and percentage of sequences are noted.

## Supplementary table captions

**Table 1-Source Data1.** ChIP-seq read counts in peak regions in replicates of MTD and RNAi samples (DiffBind analysis).

Page 1. ZLD ChIP-seq in *clamp-i* vs. MTD in 0-2hr embryos;

Page 2. ZLD ChIP-seq in *clamp-i* vs. MTD in 2-4hr embryos.

Page 3. CLAMP ChIP-seq in *zld*-i vs. MTD in 0-2hr embryos;

Page 4. CLAMP ChIP-seq in *zld-i* vs. MTD in 2-4hr embryos;

**Figure 2-supplementary table 1.** ATAC-seq read counts in peak region in replicates of MTD and RNAi samples (DiffBind analysis).

Page 1. *clamp*-i vs. MTD in 0-2hr embryos.

Page 2. *clamp*-i vs. MTD in 2-4hr embryos.

**Table 2-Source Data1.** Peaks locations in each CLAMP or ZLD-related category.

Page 1 Type I (n=5): both DA, CLAMP ZLD co-bound

Page 2 Type II (n=23): CLAMP DA and ZLD non-DA, CLAMP ZLD co-bound

Page 3 Type III (n=88): ZLD DA and CLAMP non-DA, CLAMP ZLD co-bound

Page 4 Type IV (n=434): both non-DA, CLAMP ZLD co-bound

Page 5 DA with CLAMP 0-2hr;

Page 6 DA without CLAMP 0-2hr;

Page 7 Non-DA with CLAMP 0-2hr;

Page 8 Non-DA without CLAMP 0-2hr;

Page 9 DA with ZLD NC14+12 min;

Page 10 DA without ZLD NC14+12 min;

Page 11 Non-DA with ZLD NC14+12 min;

Page 12 Non-DA, without ZLD NC14+12 min.

## Notes

### Competing Interest Statement

The authors have declared no competing interest.

### Summary of Updates

Here is a summary of the significant revisions we have made to the manuscript: 1) We re-analyzed our ATAC-seq data using a new FDR cutoff and defined new classes of loci bound by both CLAMP and Zelda to define their relative functions. We determined that CLAMP regulates Zelda recruitment by enhancing chromatin accessibility and that both factors regulate each other's target genes. 2) We discovered that the targeted role of maternal CLAMP at genes that often encode key early transcription factors, including those within the Hox cluster, is required for zygotic genome activation and viability. However, we have now determined that CLAMP acts directly in a non-redundant manner to regulate chromatin at fewer sites than Zelda does. Therefore, we have rewritten the manuscript to emphasize that CLAMP is a pioneer-like factor. 3) We added a comparison with recently published GAF ATAC-seq data (Gaskill et al., 2021 eLife) which revealed that even though GAF and CLAMP have similar binding sites, the regions at which either factor alone alters chromatin accessibility are different. 4) We defined four different subclasses of sites that are bound by both CLAMP and Zelda but are differentially regulated. We identified sites that are altered when a single transcription factor is depleted, sites where both factors are required, and a group of sites where neither factor is required. Interestingly, CLAMP and Zelda promote each other's occupancy at sites where neither factor is individually required, but both motifs are present. Therefore, it is possible that other factors such as GAF are required and/or CLAMP and Zelda are functionally redundant at a subset of sites where they are not individually required. 5) We extensively revised the manuscript to remove any potential overstatements, clarify points of confusion, and added suggested citations.

